# Engineered bacterial swarm patterns as spatial records of environmental inputs

**DOI:** 10.1101/2022.01.20.477106

**Authors:** Anjali Doshi, Marian Shaw, Ruxandra Tonea, Soonhee Moon, Anish Doshi, Andrew Laine, Jia Guo, Tal Danino

**Author notes:** Correspondence should be addressed to T.D.

## Abstract

A diverse array of bacteria species naturally self-organize into durable macroscale patterns on solid surfaces via swarming motility—a highly coordinated, rapid movement of bacteria powered by flagella^1–5^. Engineering swarming behaviors is an untapped opportunity to increase the scale and robustness of coordinated synthetic microbial systems. Here we engineer *Proteus mirabilis*, which natively forms centimeter-scale bullseye patterns on solid agar through swarming, to “write” external inputs into a visible spatial record. Specifically, we engineer tunable expression of swarming-related genes that accordingly modify pattern features, and develop quantitative approaches to decode input conditions. Next, we develop a two-input system that modulates two swarming-related genes simultaneously, and show the resulting patterns can be interpreted using a deep learning classification model. Lastly, we show a growing colony can record dynamic environmental changes, which can be decoded from endpoint images using a segmentation model. This work creates an approach for building a macroscale bacterial recorder and expands the framework for engineering emergent microbial behaviors.

## Main Text

Swarming behaviors are ubiquitously found in natural systems, ranging from bird flocks to microbial communities, and have inspired creation of artificial systems such as robot swarms^6–8^. A collective movement stemming from individual interactions, swarming can greatly increase a community’s scale as well as robustness to noisy individuals and environments. The swarming of many microbial species creates complex emergent patterns at the centimeter-scale on solid surfaces^9,10^. While a long-standing goal of synthetic biology has been to program self-organization in such a fashion, swarming motility has yet to be engineered or used for biotechnological applications^11–13^. Previous approaches have focused on prototypical microbes such as *E. coli*, which forms homogenous colonies, and have engineered swimming and quorum-sensing systems in liquid-agar environments, or utilized external pre-patterning to generate coordinated behavior^14–16^. One promising application of engineering natural swarming is the creation of a durable spatial recording system, using the sensing capabilities of millions of individual bacteria within a swarm to visibly “write” information onto a solid surface. Thus far, synthetic cellular information recording efforts have achieved recording of multiple inputs, cellular lineage, and transient signals, primarily within DNA, but rely on sequencing and other technologies for decoding^17–21^.

We focused on engineering the unique swarming of *Proteus mirabilis*—a commensal gut bacterium also commonly found in soil and water, which produces a bullseye pattern on solid agar defined by concentric rings of high bacteria density that are visible to the naked eye^22^ (**Fig. 1a**). The inherent clock-like timing and internal consistency of its ring formation naturally suggest application as a recording system, similar to the way a growing tree records information in the rings in its trunk^23^. Although the ability of *P. mirabilis* to produce rings has been known for over 100 years, it has not been developed as a synthetic biology platform, and quantification of its macroscale patterns has been limited^24^. Beyond the large-scale features of *P. mirabilis* that enable simple decoding visually, applying methods of deep learning and image segmentation can further decode multiple external inputs and dynamic conditions from more complex pattern features.

**Figure 1:**
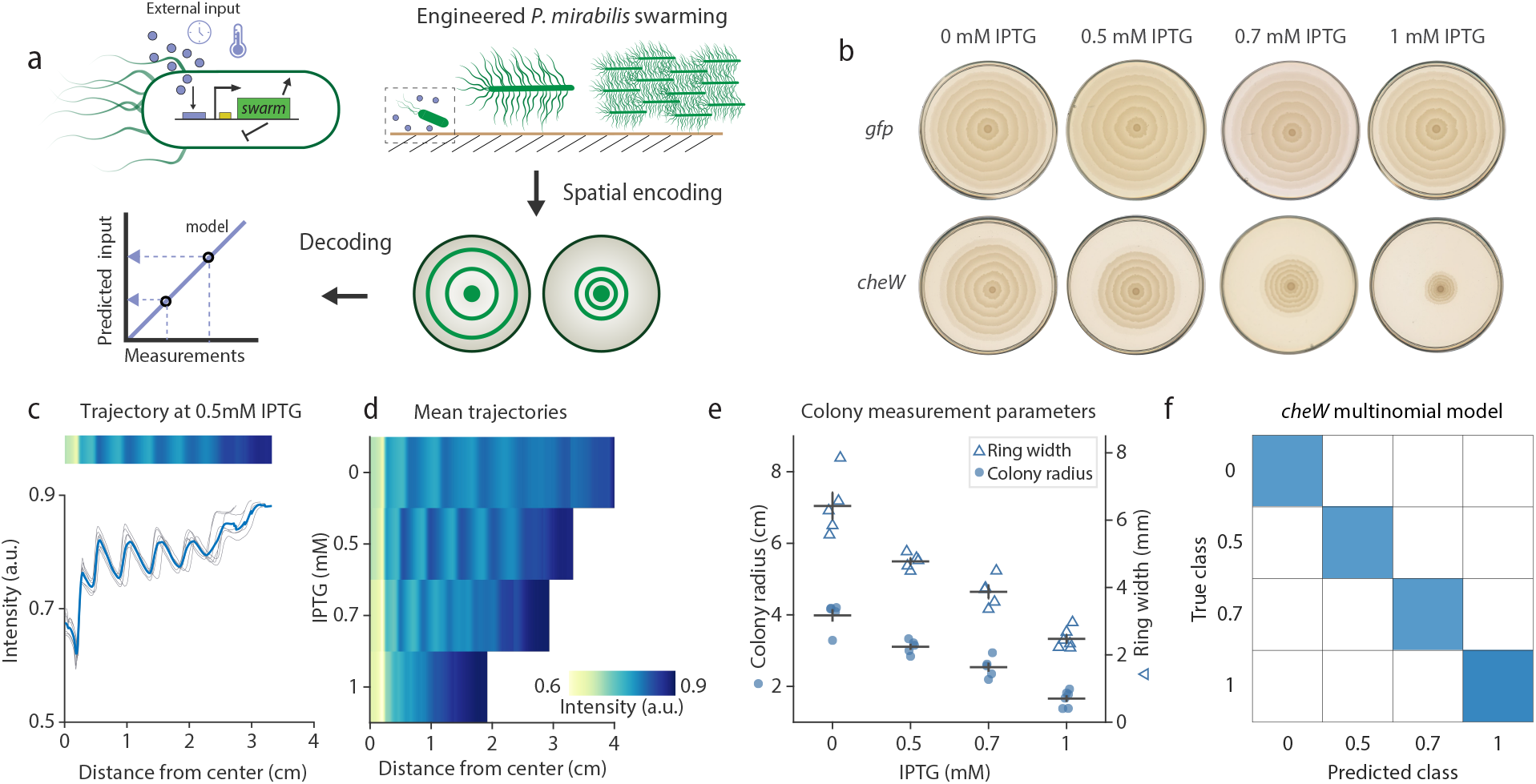
Engineered *P. mirabilis* swarm patterns as spatiotemporal records. **a.** Wild type *P. mirabilis* cells undergo oscillatory swarming on solid agar to grow into a characteristic bullseye colony via elongation, hyperflagellation, and raft formation. *P. mirabilis* is engineered with an externally inducible genetic circuit driving swarming-related genes to modify the macroscale pattern output, which can then be decoded using quantitative methods to predict the input conditions. **b.** Representative images of colony patterns formed by a strain containing a control circuit with green fluorescent protein (*gfp*) (top) compared to a circuit with the chemotaxis gene *cheW* (bottom), grown for 24 hours on agar supplemented with various IPTG concentrations. **c.** The *cheW* colony pattern is distilled into radially averaged pixel intensity profiles, with distinct peaks matching low-density ring boundaries when plotted as a heatmap or line plot. The blue line denotes the mean profile of the individual plates (each gray line represents one plate). **d.** Heatmaps of average *cheW* profiles at varying IPTG concentration (n = 5 plates at each condition except 1 mM IPTG (n = 6)). Colormap is on same scale for (**c**) and (**d**). **e.** Radii of the colonies plotted by IPTG concentration after 24 hours (filled circles) and calculated ring width (empty triangles), derived from Fourier analysis of the radially averaged profiles of individual images. The mean and standard error of the mean (SEM) are shown in black. **f**. A multinomial model was fit to the measurements in (**e**), with predicted IPTG concentration as the output variable. The model’s predictions for each plate shown in (**e**) are shown as a confusion matrix. Color reflects n per square (same as listed in (**d**); white squares represent 0).

The bullseye pattern of *P. mirabilis* is created from a sequence of phases starting with initial colony growth (lag), followed by oscillatory cycles of synchronized colony expansion (swarming), and stationary periods of cell division (consolidation). The synchronicity of its swarming is achieved by complex coordination of cell elongation, secretion of surfactant to aid movement, intercellular communication, and alignment of swarmer cells into rafts by intercellular bundling of overexpressed flagella^25, 26^. While investigation of the mechanisms governing these behaviors is ongoing, studies have identified an array of genes upregulated during consolidation phases, including those responsible for synthesis of flagella, metabolism, and cell division, and during swarming phases, such as the master regulator *flhDC*^26–30^. These works have shown that modification of expression of these genes and choice of growth conditions can lead to different variations of the ring pattern^31, 32^. We therefore chose the strain PM7002, with baseline conditions that would create a pattern with distinct ring boundaries, such that modifications to the pattern would be easily visible and quantifiable (**Fig. S1, S2**). After establishing these conditions, we expressed swarming-related genes to controllably modify specific colony pattern features, which could be subsequently analyzed and decoded to report on conditions during colony growth (**Fig. 1a**).

To initially demonstrate swarm pattern modulation, we engineered *P. mirabilis* with a high-copy plasmid carrying an isopropyl ß-D-1-thiogalactopyranoside (IPTG) inducible promoter, pLac, expressing *cheW*, a chemotaxis-related gene upregulated in the swarming process (**Fig. S12**)^29, 33, 34^. In *E. coli* chemotaxis, CheW is a membrane-bound coupling protein part of a signaling complex in which it bridges the kinase CheA to chemoreceptors, allowing phosphotransfer to CheY and CheB, where CheY is involved in control of flagellar motor rotation^35^. Although the exact role of *cheW* in swarming is not fully known, a *cheW* mutant of *P. mirabilis* was previously found to be unable to swarm^33^. Here, inducing constitutive *cheW* expression with increasing concentrations of IPTG in the agar generated colonies of decreasing ring width and size at 24 hours, compared to a control *gfp*-expressing strain which showed no change in pattern in response to IPTG (**Fig. 1b**). To quantify the patterns, we examined the radially averaged pixel intensity as a proxy for colony density in each of the conditions, where high pixel intensity (light colors) represents lower density (**Fig. 1c, Fig. S3**). All colonies had a characteristically dense boundary around the central inoculum, seen as a dip in the intensity plot around 0.25 cm from the center of the colony (x=0 in the plot), and showed periodic changes in density across the colony. As expected, the radially averaged intensity profiles showed peaks of intensity corresponding to the periodic ring boundaries. With increasing *cheW* expression, the profiles showed greater density near the inoculum and at the ring boundaries, which can be seen as lighter areas on heatmaps of radially averaged intensities (**Fig. 1d**). We constructed a small dataset and measured colony radius manually using image processing tools, and ring widths using a custom algorithm (Methods) (**Fig. 1e**). The colony radius and ring width correlated well with IPTG concentrations (R^2^= 0.90 for each). To more accurately decode input conditions from the pattern, we fit a multinomial regression model on these measurements and found that the model correctly predicted each colony’s input IPTG from the combination of its radius and ring width in all cases (**Fig. 1f**). We thus reasoned that colony features could potentially encode information about external inputs received by the bacteria, and feature measurements could subsequently be used to decode the information.

### Manipulation of multiple swarming-pathway genes

Given the variety of features observed in *P. mirabilis* patterns in the literature beyond the ring widths and overall colony radii, we explored the potential for multiplexed information encoding. Here we sought to identify additional genes which could distinctly modulate colony pattern features (**Fig. 2a**). We chose genes previously implicated in a range of points in the swarm process, including *umoD*, which controls the master regulator of swarming, *flhDC*; the signaling factors *fliA* and *flgM*, which are involved in flagellar gene transcription; and *lrp*, which affects general cellular processes in response to leucine presence^29, 36–40^. Induced expression of these genes via IPTG generated a variety of patterns, ranging from dense ruffled textures, to “spikes”, to indistinct ring boundaries (**Fig. 2b**). Scanning a range of IPTG concentrations showed graded changes in patterns (**Fig. S4)**. For example, with minimal IPTG-induced expression, the *lrp* strain formed spikes in the inner colony rings, and at maximal induction each ring boundary was spiky. Increased expression of *flgM* caused colony radius at 24 hours to shrink, while *fliA* caused the formation of more visible dots or “microcolonies” just within the boundaries of each ring. As *umoD* expression increased, colonies became more symmetric, and ring boundaries and the inoculum edge became fainter. Taken together, these various qualitative characteristics suggested that induced expression of certain swarm genes could indeed affect several pattern features which could be measured and quantified.

**Figure 2:**
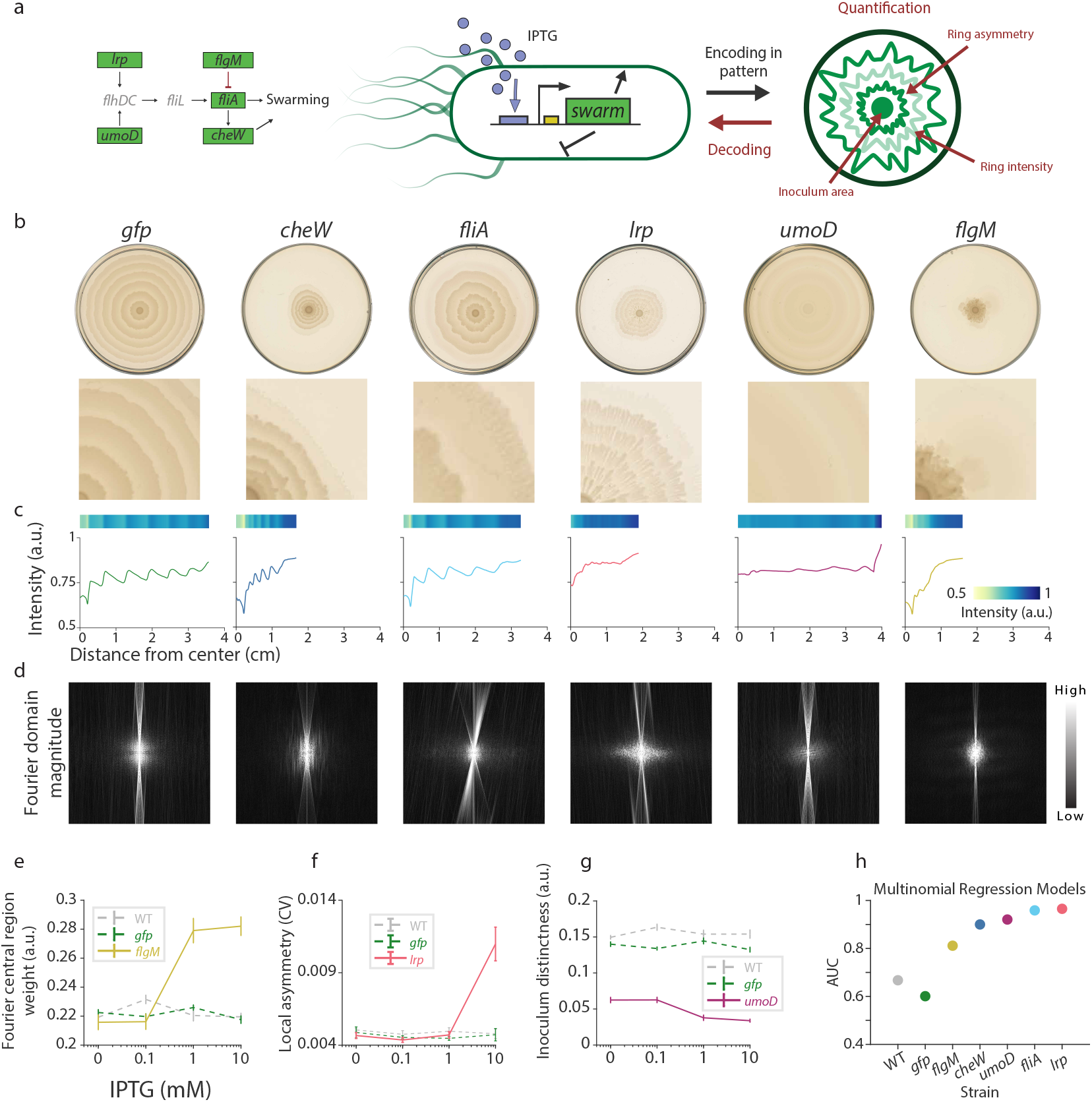
Modulation of swarm genes in engineered *P. mirabilis* results in quantifiable changes to distinct spatial pattern features. **a.** Candidate genes involved in *P. mirabilis* swarming pathway were chosen for construction of inducible strains. The patterns in the presence of inducer were characterized by growth on IPTG-supplemented solid agar, then by specific feature measurements used to recover the inducer concentration. **b.** Characteristic patterns of engineered strains in the presence of IPTG and closeups of pattern features. **c.** For each induced strain, heatmaps and plots of radially averaged intensity profiles across the colony for the representative images in (**b**). **d.** Fourier transforms of the polar images visualize the magnitudes of the intensity frequencies of each induced strain. **e-g.** Quantification of aspects of colony patterns of engineered strains at increasing IPTG concentrations. All strains had at least n = 3 plates measured at each IPTG concentration. Error bars represent standard error of the mean (SEM). Details can be found in Methods. **e.** Intensity of central region compared to total intensity of the Fourier transform of the polar image. **f.** Local radial coefficient of variation (CV), which increases with colony asymmetry. **g.** Change in intensity from the densest edge of the inoculum (innermost circular region of colony) to the low-density region immediately surrounding it, i.e., distinctness of the inoculum edge, where low values correspond to less distinct edges. **h.** Area under the curve (AUC) of multinomial regression models for predicting IPTG concentration, fit with specific pattern measurements for each strain.

We next examined the radially averaged profiles of each pattern, which revealed distinct characteristics for each strain (**Fig. 2c**). For example, overall colony density was higher with induced *cheW* expression than with *umoD*. The spikes visible in the *lrp* pattern, which caused ring boundaries to spread over greater widths, reduced the sharpness of the ring boundaries in the radially averaged profiles. Given the repeating nature of features in the patterns, we also explored visualization of the Fourier spectra of the polar transforms of the images, which highlight the presence of frequency information, to see if the spectra varied between strains (**Fig. 2d**). The periodic features in the patterns resulted in high visible intensities in certain regions of the Fourier transform images. For example, the *umoD* strain displayed a higher magnitude in the outer regions of the transformed images, representing high-frequency (i.e., short distances between repeated features) information, while the other strains showed greater magnitude at the central regions, which represents lower-frequency features. In summary, we saw that the visible differences in patterns between the engineered strains were reflected in, and thus could be analyzed from, quantitative representations of the images.

We next sought to identify features of each strain’s pattern which could allow for determination of the input IPTG concentration. We generated a dataset of images for each strain grown at a range of inducer concentrations and measured a range of features for each (**Fig. S5a**). The low frequencies of the Fourier spectra were found to increase with IPTG induction for the *flgM* and *cheW* strains, reflecting the visual observation of thinner, fewer rings of increased density at higher IPTG. (**Fig. 2e**). A second measure, the local coefficient of variation (CV), increased with increasing IPTG for the *lrp* strain, which could be observed visually in the spiked rings (**Fig. 2f**). Finally, the distinctness of the inoculum border, measured by the change in intensity over the border, decreased with increasing IPTG for the *umoD* strain, particularly from 0.1 to 1 mM IPTG. (**Fig. 2g**). These measurements showed that induced expression of these genes could quantifiably affect the pattern in response to changes in IPTG.

As an approach for decoding information from the patterns, we explored fitting regression models on these measurements. The samples were binned into three classes (0-0.09, 0.1-0.9, and 1-10 mM IPTG), and then each feature individually, and all possible combinations of the measured features, were used to fit multinomial regression models, to identify which combination would best decode a given strain’s pattern. The performance of such models can be evaluated using a multi-class area under the receiver-operating curve (AUC) metric, where the more accurate a model is for predicting true positives compared to false positives for each class, the closer the AUC will be to 1. The AUC of each strain’s fitted model was evaluated on the input data (**Fig. 2h, S5b**). For each strain, the combination of parameters which gave the highest AUC varied, confirming that each strain was encoding information in a characteristic combination of pattern features. The best models for the experimental strains with *cheW*, *fliA*, *lrp* generally had AUC>0.9, showing that the models were well able to differentiate true positives in each IPTG class from false positives. The AUCs were 0.6 for the *gfp* control strain, just slightly above a random classifier (AUC=0.5), suggesting that pattern parameters were not strongly affected by increasing IPTG for control strains. The confusion matrices showed that the fitted models correctly classified a majority of the plates for each strain (**Fig. S5c**). Thus, information about the environment encoded within the engineered strains’ patterns can be decoded using combinations of relevant pattern features.

### Dynamics of engineered *P. mirabilis* strains

*P. mirabilis* swarming creates patterns not only in space, but also in time; this temporal regularity suggests the possibility of encoding information in both the endpoint patterns and their dynamic growth phases. We aimed to gain an understanding of the dynamics of the engineered strains by time-lapse imaging of colony growth (**Fig. 3a**). In order to capture high-resolution images of swarming, we developed a time-lapse setup using a commercial flatbed scanner. For each strain, a time-lapse was captured with maximal IPTG concentration at 25°C; images were taken every 10 minutes over the course of the time-lapse (**Fig. 3a, Movie S1**). The individual images were then radially averaged and full time-lapses were visualized via heatmaps (**Fig. 3b**). Using a custom semi-automated algorithm (see Methods), we identified the location of the colony front at each timepoint and obtained trajectories with high spatiotemporal resolution (**Fig. 3c**). The colony growth trajectories showed that each of the engineered strains maintained the classic alternation in phases, but with changes in aspects such as initial lag time and length of the phases compared to the control *gfp* strain. We then measured the mean length of time of each phase from each of these trajectories (**Fig. 3d**), which, together with distance swarmed during each swarm phase, enabled the calculation of swarm speed (**Fig. 3e**).

**Figure 3:**
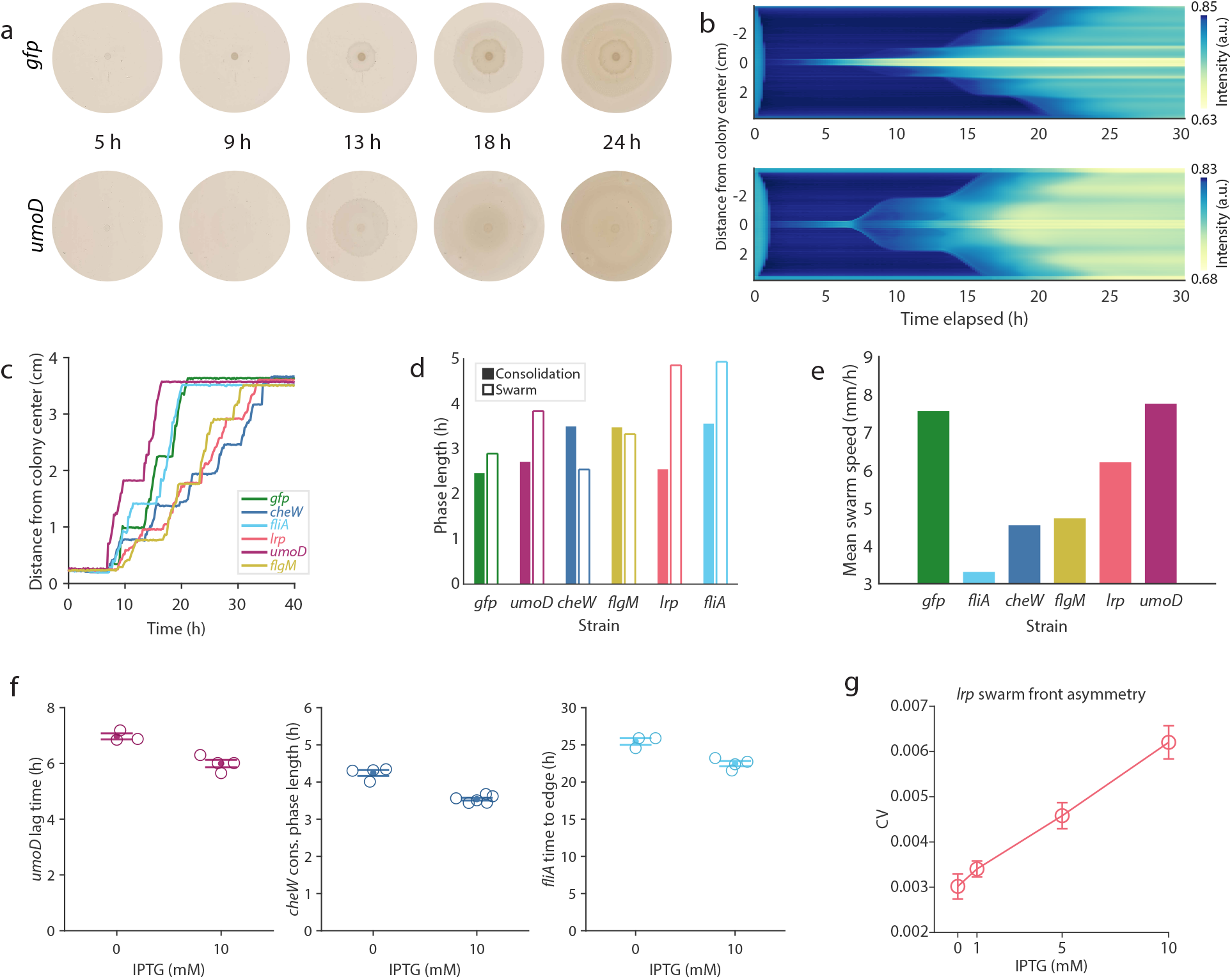
Dynamics of engineered *P. mirabilis* pattern formation. **a.** Time-lapse of *P. mirabilis* with inducible *umoD* expression or inducible *gfp* expression (control). Plates contained 20 mL 1.3% agar with 10 mM IPTG. **b.** Heatmap visualizations of swarming pattern development from center of plates (0 cm on left axis) to edge (top and bottom edges) for each image in the time-lapses in (**a)**. Radially averaged pixel intensity, a proxy for local colony density, at each location on plate is represented by heatmap color, with blue indicating least dense and yellow indicating most dense regions. Active regions and time periods of colony expansion via swarming are visible as faint blue diagonal edges. Consolidation phases appear as horizontal edges corresponding with increasing density (lighter colors) within the colony. **c.** Colony front distance from center plotted as a function of time for a single time-lapse of six plates. All plates contained 10 mM IPTG. **d.** Mean consolidation (filled bars, left) and swarm (outlined bars, right) phase lengths calculated from the trajectories in (**d**). **e.** Mean of the swarm speeds for each strain in the same time-lapse. **f.** Measurements of dynamic features at 0 vs 10 mM IPTG for the indicated strains. Each condition and strain was tested on at least n = 3 separate plates. All plots represent a significant difference between induced and uninduced conditions (p-values from a 2-sample t-test were 0.003, 2e-5, 0.003 for the plots of *umoD*, *cheW*, and *fliA* respectively.) **g.** The local CV of the swarm front for a colony (averaged over all swarm phases each colony underwent, 3 phases at 0 and 1 and 4 phases at 5 and 10 mM IPTG) at each given IPTG. Error bars in (**f**) and (**g**) represent SEM.

To explore whether certain dynamic parameters would show a trend with increasing IPTG for each strain, we generated individual time-lapses of each strain grown at a range of IPTG concentrations (**Figs. S6-7**). When comparing uninduced to induced conditions, we observed distinct measurements for each strain such as the lag time for *umoD*, the length of the middle consolidation phases for *cheW*, and the time for the colony to cover the plate for *fliA* (**Fig. 3f**). More complex dynamic parameters also encoded information; for example, the asymmetry of the colony front during swarming phases increased with IPTG for the *lrp* strain (**Fig. 3h**). These results suggest that dynamic parameters can also be used to encode and decode information from these spatiotemporal patterns, and that in the future strains can be chosen for a given application depending on the desired time scale of recording.

### Multiplexed recording using a dual-input strain

In order to build a strain which could provide information about multiple inputs simultaneously, we induced a second swarming-related gene with the pBAD operon and promoter, transcribed in the presence of arabinose (**Fig. S9**). Since swarming-related genes have interdependent effects, we sought to try two genes which robustly changed distinct pattern features on their own. We thus built a combination strain with *cheW* expression induced by the pLac promoter, and *umoD* expression induced by pBAD promoter (**Fig. 4a, Fig. S9**). Initial characterization of this strain demonstrated that its swarm patterns indeed distinctly reflected the presence or absence of each input (**Fig. 4b**). Representative radially-averaged profiles were visualized as heatmaps for comparison (**Fig. 4c**). The plates imaged followed a characteristic pattern at most of the conditions. Increasing IPTG from 0 to 1 mM, inducing *cheW* expression, resulted in a visible decrease in 24-hour colony radius, ring width, and colony symmetry, as seen previously in the single input strain. Meanwhile, increasing arabinose from 0 to 0.1% resulted in a highly symmetric pattern with initially semi-distinct, narrow rings giving way to the indistinct wide rings more characteristic of the single-input *umoD* pattern. The combination of IPTG and arabinose presence resulted in a similar pattern, with narrower inner rings giving way to wider outer rings, but with smaller colonies at 24 hours and asymmetric ring boundaries compared to those formed with arabinose alone.

**Figure 4:**
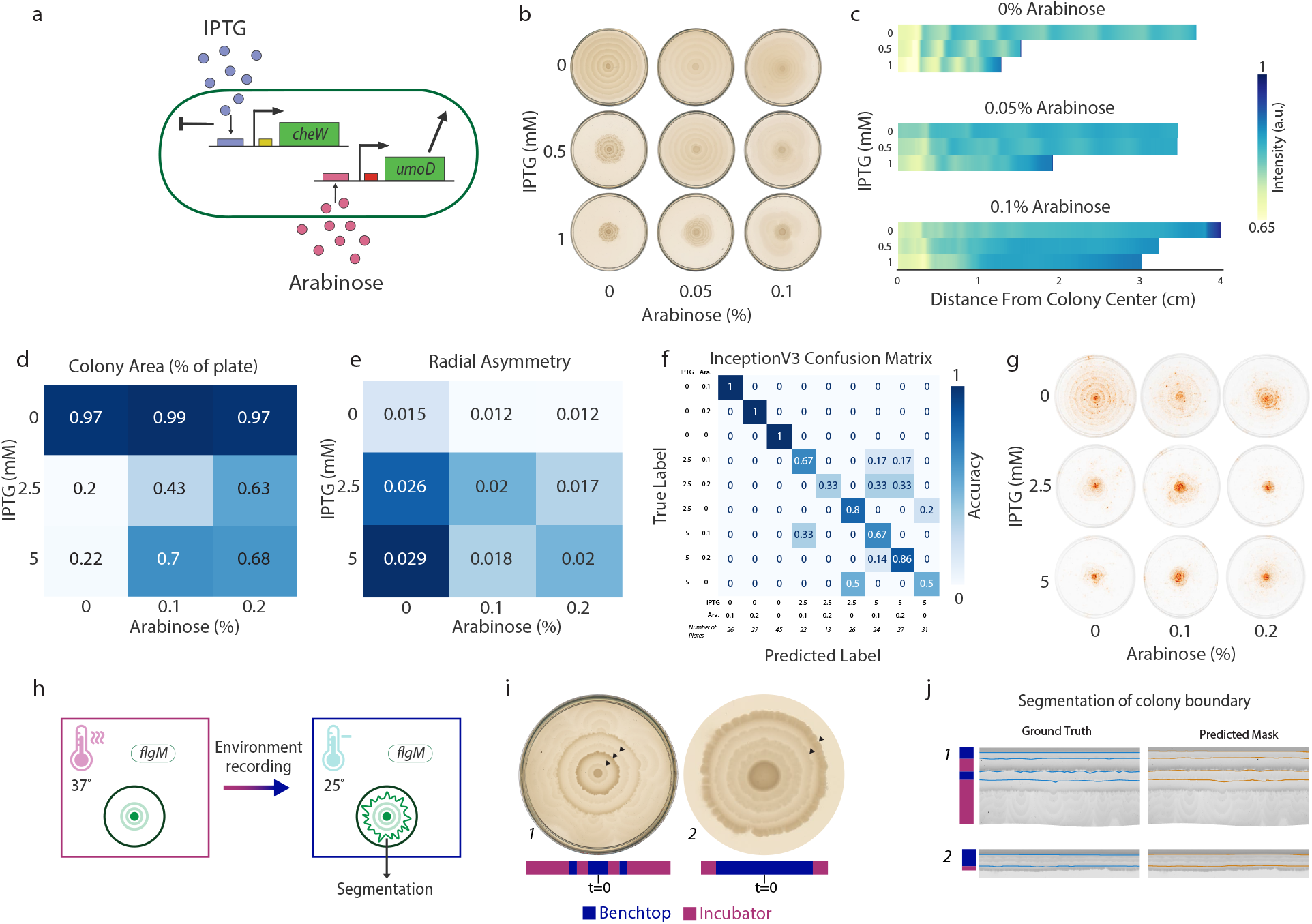
Multi-condition pattern encoding and deep-learning models for decoding. **a.** Dual input swarming strain with IPTG-inducible expression of *cheW* and arabinose-inducible expression of *umoD*. **b.** Representative images of colony patterns produced by the dual *cheW/umoD* strain in response to combinations of IPTG and arabinose. **c.** Heatmaps of radially averaged profiles of the patterns in (**b**) are shown. **d-e.** Mean colony area (calculated as percent of agar area in flattened image) and coefficient of variation for all plates (n>=14) at each combination of IPTG and arabinose. **f.** Confusion matrix for the InceptionV3 model’s accuracy of predicting combinations of IPTG and arabinose concentrations from endpoint patterns unseen during training. Total available images per class shown below matrix; an 80/20 train/test split was used. Numbers on matrix represent fraction of test images per true class. **g.** Visualization of the pixel attributions from the InceptionV3 model for representative, correctly-predicted images of each class. Darker orange represents higher weight of that pixel on the final prediction. **h.** Schematic of encoding of environmental changes within developing *flgM* pattern. **i.** Example patterns of the *flgM* strain grown with 10 mM IPTG and moved between the benchtop and incubator. Arrows mark boundaries between regions of the pattern formed in different conditions, i.e., the location of the colony edge at the time of a switch in conditions. **j.** Examples of the predicted boundary masks generated by the trained U-Net compared with ground truth annotations for the pattern images shown in (**i**), which were unseen during training.

To characterize the *cheW* and *umoD* combination patterns in more detail, a dataset of plate images at IPTG concentrations of 0, 2.5, and 5 mM combined with arabinose at 0%, 0.1%, and 0.2% was created. The average percent of the plate covered by the colony at each condition decreased with increasing IPTG and increased with the addition of arabinose (**Fig. 4d**). However, increase of arabinose from 0.1% to 0.2% had little effect on the colony area except at 2.5 mM IPTG (**Fig. S9**). Similarly, average radial CV as a measure of colony asymmetry increased with the induced expression of *cheW*, but decreased with the addition of arabinose inducing *umoD* expression (**Fig. 4e, S9**).

As done previously for the single-input strains, a set of standard measurements was then taken on each image in the dataset, and a 9-class multinomial regression model was fit on the output (**Fig. S10**). The model performed poorly, predicting almost all images as 0% arabinose, and the maximum AUC achieved was only 0.72. This result suggested that the two-input strain’s patterns, involving interdependent swarm genes, were too complex for the previous regression-based decoding method. However, the ease of distinguishing the patterns by human eye suggested that the application of deep learning methods for image classification could prove useful for decoding the patterns. In particular, deep convolutional neural networks (CNNs) have clear applicability and have not yet been used to characterize macroscale bacterial colony patterns. CNNs can learn to extract salient features from bacterial images and classify patterns to predict the image class^41^.

We fine-tuned models including ResNet and the Google Inception V3 networks to classify images in the dataset into one of the nine classes (details in Methods). The models were pre-trained on ImageNet data, a common strategy in deep learning (**Fig. S11**)^42^. Here, the fine-tuned Google InceptionV3 model was able to successfully classify the majority of our images (**Fig. 4f**). An ROC curve was calculated (see Methods) and the AUC was 0.96, a noticeable improvement from the multinomial regression model. Such models can also be characterized by “top-3” accuracy, i.e., when used to predict the three most likely classes of an image, whether one of the three is the correct class; the fine-tuned model achieved a top-3 accuracy of 0.98. We observed that intermediate concentrations of IPTG and arabinose reduced the model’s accuracy due to some bimodality in pattern formation (**Fig. 4f**). Visualizing the pixel attributions of the model indicated the inoculum and inner rings had a large impact on the predictions, suggesting that these areas of the pattern were most affected by the induced expression of the different swarm genes (**Fig. 4g**). Since the innermost portion of the colony was most critical to pattern prediction, pattern decoding may be possible after just a few hours of growth, rather than needing to wait 24 hours until the full plate is covered. Overall, these results suggest that our system can be used to encode and decode multiple inputs, and that the use of deep networks along with transfer learning will enable decoding of complex pattern feature changes.

### Multi-condition pattern segmentation and information decoding with deep learning

We next sought to determine whether an engineered strain could record changes in the environment taking place during pattern formation and how these changes could be decoded from the endpoint pattern, similar to the analysis of rings in a tree^23^. We used the *flgM* strain, which we had observed to form two strikingly different patterns in the presence of 10 mM IPTG in the incubator vs on the benchtop: a swarming-inhibited, ruffled, dense pattern in the incubator at 37°C, and a wide-ring, symmetric, less dense pattern on the benchtop at ~25°C (**Fig. S11d**). After inoculation, plates were first placed in one condition; after some time, plates were switched to a second condition, and certain plates were switched a third time before the endpoint scans were captured (**Fig. 4h**). Plates were scanned before each switch, creating a dataset of 21 images. Representative pattern images are shown in **Fig. 4i.** This shift in environmental conditions resulted in the formation of rings alternating between indistinct, radially symmetric, wider rings and dense, asymmetrical, narrow rings, visible as bands on the polar-transformed images (**Fig. 4j**). In general, denser regions corresponded to incubator growth, while fainter regions with wider rings corresponded to benchtop growth.

To decode these alternating ring patterns, we manually annotated the dataset, creating ground truth masks of the boundaries marking the shift in the pattern corresponding to a shift in the environment. We then trained a U-Net model, a type of network frequently used for segmentation problems, pretrained on ImageNet to predict these boundaries given an input pattern image (details in Methods). Our model achieved above 95% training and validation accuracy and above 90% recall within the first 25 epochs of training, showing that it could learn the features within the dataset (**Fig. S11e**). Application of the trained model to previously unseen images resulted in specific prediction of boundaries matching the ground truth, and noticeably did not simply highlight all ring boundaries (**Fig. 4j**). In future, these predicted boundaries could be used to back-calculate the time at which a given perturbation was experienced, by generating prior control measurements of the time of formation of rings at different conditions. Taken together, these results demonstrated that our approach could be used to decode information about changing environment from the engineered strains’ patterns.

## Discussion

We have developed a proof-of-concept approach to engineering spatial patterns in *P. mirabilis* for information encoding and decoding. To date, bottom-up efforts to control spatiotemporal behaviors in microbial synthetic biology have required complex genetic circuits, used *E. coli* strains with liquid media, or required externally pre-patterned cues^43–45^. While there have been recent advances in encoding information in DNA and fluorescent bacterial colonies, there has not yet been an attempt to apply macroscopic pattern engineering for encoding information^17, 21, 46^. The approach described here takes advantage of the natural pattern formation capabilities of *P. mirabilis* on solid agar coupled with synthetic biological engineering approaches to modulate durable swarm patterns. We constructed genetic circuit variants with swarming-related genes and developed automated approaches for decoding information by quantifying aspects beyond typical colony radius measurements, such as colony asymmetry, swarming speed, frequency spectrum, and inoculum border distinctness. We then expanded to a dual-input system to sense two inducers, and trained a deep learning classifier to decode its patterns; while some works have begun to apply deep learning for segmentation of macroscale colonies, and in several cases for microscopic cell segmentation or classification of smaller colonies, our work represents a new application for classification, that of complex macroscale colony patterns ^47–51^. At the same time, the macroscale patterns had many attributes distinguishable by eye, which could enhance the applicability of this system.

Since external conditions do affect pattern formation, a practical consideration for use of this system is to reliably produce robust patterns in differing laboratory or field conditions. We envision the use of this platform with side-by-side controls not exposed to the environment or input of interest, such that relative differences in pattern changes could be recorded. Additionally, future versions of the current system could include construction of knockout strains as well as chromosomal integration of promoter systems, which may allow for tighter control over the final pattern. In particular, for the dual-input strain, at the intermediate condition (2.5 mM IPTG and 0.1% arabinose), two distinct groups of patterns emerged, one in which colonies were small and dense, and one in which colonies swarmed almost to the edge of the plate. This stochasticity could possibly be reduced in future with further engineering, a different combination of genes or a different range of concentrations of inputs, which in turn can allow the decoding models to achieve higher accuracy. Enhanced imaging approaches such as incorporating a pigment into to the swarm medium or using pigment-producing strains may also improve accuracy. Further development of algorithms for image processing will benefit from the training and application of deep learning models for segmentation of colony and ring boundaries, such as the pipeline we have recently developed^48, 52^. Additionally, the application of increasingly sophisticated computational approaches for modeling and machine learning-based classification will allow for the use of more complex spatiotemporal patterns^48^. Such models can be incorporated into easy-to-run computer or mobile applications, and optimized for use with cell phone camera-images, allowing on-the-go analysis with inexpensive technologies. While we aimed to standardize our data acquisition method so that lighting, image size, and other factors would be constant throughout the datasets, these aspects can be intentionally varied to capture a more diverse dataset, which could help in developing models for application in a broader range of settings.

Beyond these improvements, the proof-of-concept system presented here can be expanded in several directions. The approach could be used to explore other inputs such as light, radiation, or gaseous molecules, or to develop a longer running recorder for changes in temperature or air quality. Other swarming species with natural swarming properties could be manipulated such as *Pseudomonas aeruginosa*, *Paenibacillus vortex*, or *Bacillus subtilis ^2, 3, 53^*. Controlling swarming behaviors by engineering bacteria can enable multiple applications, ranging from bacteria drug delivery to living material assembly. The approaches developed here can in turn shed light on *P. mirabilis* growth dynamics and virulence, and be applied to understanding the coordinated and emergent behaviors of microbes.

## Methods

### Bacterial strains and growth conditions

*Proteus mirabilis* (ATCC 7002) was kindly provided by Dr. Philip Rather. *Escherichia coli* Mach1 for cloning was purchased from Fisher. *P. mirabilis* and *E. coli* were cultured in Luria-Bertani (LB) media (Sigma-Aldrich) supplemented with 50 μg ml^-1^ kanamycin, respectively. *P. mirabilis* was grown on either 3% or 1.5% agar to suppress or allow for swarming, except for time-lapse assays as indicated.

### Competent cell preparation

*P. mirabilis* (PM7002) cells and *E. coli* (Mach1) were made electrocompetent as follows. A fresh 2-mL overnight culture was subcultured 1:100 in 50 mL LB media, then grown at 30°C with shaking until logarithmic growth phase was reached, indicated when the optical density at 600 nm (OD_600_) was 0.4-0.6. Growth was stopped by incubation of the culture on ice for 15 minutes. Cells were then pelleted by centrifuging for 10 minutes at 4°C and 3000 rpm. After decanting, the pellet was washed three times in either 50 mL ice-cold filter-sterilized 10% glycerol (*P. mirabilis*) or 50 mL ice-cold filter-sterilized water (*E. coli*), then resuspended in 220 μL 10% glycerol. 50 μL aliquots were stored in −80°C.

### Strain construction

The previously constructed pZE24 (pLacGFP pConstLacIQ) plasmid, containing the ColE1 origin of replication and a kanamycin resistance cassette, was used as the backbone for the inducible swarming plasmids. Plasmids and chromosomal *P. mirabilis* DNA were prepared using standard procedures (Quiagen). Swarming gene sequences were obtained from GenBank (JOVJ00000000.1) and Gibson primers were designed (Eton) to amplify the genes from the chromosomal DNA via PCR (Phusion)^54^. A set of swarming plasmids were constructed using Gibson Assembly and standard restriction digest and ligation cloning to replace the *gfp* gene with the appropriate swarming gene. For plasmids which additionally contained *pBAD-araC*, the operon was obtained from the pBADmCherry-pConstAra plasmid (ATCC54630). After cloning plasmids into Mach1 *E. coli*, clones were verified via colony PCR (Phusion) and sequencing (Eton). Clones were then grown at 37°C with shaking overnight before being stored in 50% glycerol at −80°C. All plasmids and strains are listed in **Tables S1 and S2**; plasmid maps are shown in **Fig. S12**.

### *P. mirabilis* transformation

Plasmid DNA was introduced into *P. mirabilis* competent cells as follows. 50 μL aliquots of competent cells were thawed on ice for 10 minutes. DNA was added to the cells (200-400 ng DNA in a volume of 1-5 μL per aliquot). The mixture was then incubated on ice for one hour. Cells were electroporated in prechilled electroporation 0.1 cm electrode gap cuvettes using a Bio-Rad MicroPulser set to E1 setting (1.8 kV) for bacterial electroporation. Cells were recovered by adding 1 mL prewarmed SOC media and incubated with shaking at 37°C for 3 hours. The cells were pelleted by centrifugation for 10 minutes at 4°C and 3000 rpm, and 700 μl of the supernatant was decanted before resuspension in the remaining 300 μl. The cells were then plated on pre-warmed 3% LB agar plates with antibiotics as necessary and incubated at 37°C for 22-24 hours. Single colonies were inoculated and fresh overnight cultures were stored in 50% glycerol at −80°C.

### Bacterial growth and swarm assay

Overnight liquid bacterial cultures were prepared by inoculating LB media with cells from the −80°C glycerol stocks and supplementing with 50 μg ml^-1^ kanamycin as appropriate. Cultures were incubated at 37°C with shaking for 12-16 hours. The OD_600_ of each culture was measured and normalized to 1.0 by dilution with LB media. Swarm assays were optimized from a protocol adapted from literature. A study to develop standard conditions is shown in **Fig. S1**. Precise maintenance of the selected conditions was necessary to achieve comparable results^55^. 1.5% agar (or, where indicated, 1.3% agar) was autoclaved, then cooled to 50-55°C with stirring. 5 μg ml^-1^ kanamycin, IPTG and/or arabinose were then added as necessary. 15 mL agar was poured in each 100×15 mm Petri dish and left to solidify partially uncovered under an open flame for exactly 30 minutes. 2 μL of the previously diluted liquid culture was inoculated on the center of each Petri dish and dried for 15 minutes partially uncovered under open flame. The plates were incubated at 37°C for 24 hours, then individually imaged using a scanner (Epson Perfection V800 Photo Scanner) set to 48-bit Color and 400 dpi, with the lid off and colony side facing up. The scanner was kept on the benchtop and room lighting was similar during all experiments; other settings of the scanner were also kept constant between experiments. Incubator humidity typically varied between 50-80% during the course of experiments.

### Time-lapses

For time-lapses on the benchtop (room temperature), up to six plates with 20 mL 1.3% LB agar were inoculated and placed on the flatbed scanner using the previously described settings, and kept upside down to prevent condensation and with lids on to prevent contamination. A custom AppleScript was written to scan plates every 10 minutes for a pre-set length of time (typically 48-72 hours). Typical benchtop conditions were 25°C and 40-50% humidity.

### Computational methods

Measurements of colony features were taken using MATLAB (Mathworks) image and signal processing functions. Images were preprocessed by conversion to grayscale; the plate rim was removed using the imfindcircles (based on a Hough transform) and regionprops functions, then the image was thresholded to find the colony’s center inoculum, typically easily identified by its dark boundary. Upon finding the center point, the colony was unrolled or “flattened” using a Cartesian to polar transformation and the scattered interpolant function, and resized to 1000×1000 pixels for ease of scaling analysis for the full dataset. The colony rim was also masked out (set to white). Radial profiles could then be easily generated by averaging the pixel intensity across each row of the image. The ring widths in **Fig. 1** were calculated by using 1-D Fourier/inverse Fourier transformation on the radial profile of each image of interest to filter out noise and by subsequent peak-finding. The ring widths of a single image’s radial profile were averaged to generate the individual measurements in **Fig. 1e**.

The colors selected for plots of the different strains in figures **2-4** were derived from a previously developed “bright” color scheme^56^. Where described, *local* CV was calculated by moving a sliding window region of width 10 pixels across each row and calculating the CV within it, then taking the average of these calculated CVs over the whole image. *Mean* CV was calculated by obtaining the CV across each row, then averaging over all the rows. The inoculum edge intensity was measured for a given image as follows: the image was smoothed using the movmean function with averaging applied in 25-pixel windows horizontally. For each individual column of the smoothed image, the minimum value between the 15^th^ and 60^th^ rows (ie, in the region of the inoculum border) was subtracted from the maximum value in that region. The average over all the columns was then taken (calculation schematics in **Fig. S5a**).

For certain measurements, a mask of the colony region was desired. A custom algorithm was developed using image processing functions in MATLAB. Briefly, a set of filters were applied to reduce local noise such as dust and scratches, then adaptive histogram equalization was applied to increase contrast. The entropyfilt function in MATLAB was applied and the output was thresholded, then the difference between this output and the original image was taken in order to sharpen the edges in the colony. The image was binarized and a series of morphological operations including dilation, opening, and hole-filling were applied to obtain a mask of the colony. The largest region was retained and all smaller regions were discarded.

In order to analyze the time-lapses, a method to track a growing swarm colony was sought; such methods have been of recent interest^48, 57^. *P. mirabilis* presents a unique challenge in this area; during its swarm phase, only a thin, almost transparent film of bacteria moves outwards, almost indistinguishable from local variations in agar intensity. Thus, the swarm front is difficult to detect with conventional thresholding-based or edge-detection algorithms which have been implemented previously for analysis of other species^58–60^. The colony region isolation algorithm described above also did not work on these images. The movement outwards on the plate (or vertically down on the flattened images) over time is difficult and noisy to capture. Towards an algorithm for tracking the swarm edge, each time-lapse image was first flattened as described above. Each image was subtracted from the preceding image using the imabsdiff function. The difference images were then averaged across columns, creating a radially averaged trajectory. In brief, the findpeaks function was used on each timepoint’s trajectory, using a custom algorithm and manual parameter refinement to determine the location in which to seek the peak, and taking advantage of the constraint that the colony edge would not move backwards over time. The user could choose (1) the minimum possible prominence of the peaks and (2) the range to the right of each previous peak in which the algorithm would seek the next timepoint’s peak, and then the algorithm would iterate over the whole time-lapse. The process would be repeated until the user was satisfied with the visual overlay of identified peaks on the time-lapse heatmap. The obtained colony front trajectory was then labeled using a custom algorithm involving the moving_polyfit function, bwareaopen and bwlabel, from which the locations of the lag phase, swarm phases, and consolidation phases were obtained^61^. In **Fig. 3f**, the *cheW* measurements were calculated by discarding the first and last consolidation phases and measuring the length of only the middle consolidation phases.

Statistical tests were calculated and data was plotted either in MATLAB or in Python. Latex tables were generated using Overleaf. Multinomial regression models were fit to the measurements using the mnrfit function in MATLAB, returning the coefficients and p-values in **Fig. S5b**. For the single input strain data in **Fig. 2h**, each flattened image was divided into four sectors (each 250 pixels wide) and measurements were taken on each sector to increase the number of measurements available, so that the model fitting could converge. The models were evaluated using the multiClassAUC function, which implements the Hand and Till function for area under the curve for multi-class problems^62^. Machine learning models were implemented in Tensorflow and Pytorch, with manual annotation of the *flgM* ground truth segmentation done using the LabelMe program^63^. Attributions in **Fig. 4g** were calculated following the Integrated Gradients method of Sundarajan et al^64^. Fine tuning of the pre-trained models for classification of the dual-input strain was done with on-the-fly augmentation of the dataset, using random rotations, translations, and horizontal flips^65–67^. For the U-Net segmentation work, a VGG-11 Encoder pretrained on ImageNet was used^68–70^. Predicted masks from the U-Net model were postprocessed using standard methods. In brief, the predicted masks were dilated to ensure a given boundary was fully connected, then opened to remove any small instances of detected noise. The cleaned masks were then skeletonized to obtain single-pixel thick boundaries for evaluation of metrics such as accuracy. Finally, a flat line-shaped structuring element was applied to dilate near the left and right edges to re-connect the boundaries with these edges. For the visualization in **Fig. 4i**, masks were dilated with a disk element for better visibility.

## Acknowledgments

We thank Professor Martina Pavlicova for helpful discussion of the statistics methods used. We thank the members of the Danino lab for review of the manuscript. We thank R. Minyety for assisting in time-lapse experiments.

## Author contributions

A.D. and T.D. conceived and designed the study. A.D., M.S., R.T., and S.M. performed experiments and constructed the dataset. A.D. and M.S. performed the computational analysis. A.D., M.S., and A.D. (Berkeley) carried out the deep-learning work with input from J.G. and A.L. A.D. and T.D. wrote the original manuscript draft and A.D., M.S., and T.D. edited the manuscript with input from all authors.

## Data and materials availability

All data is available upon reasonable request. Correspondence and request for materials should be addressed to T.D.

## Code Availability

All code is proprietary and managed by the Columbia Technology Ventures Office of Intellectual Property. They are available from the corresponding author T.D. upon reasonable request, after permission from the Columbia Technology Ventures Office of Intellectual Property.

## References

1. Kearns, D.B., A field guide to bacterial swarming motility. Nature Reviews Microbiology, 2010. 8(9): p. 634.

2. Ingham, C.J. and E. Ben Jacob, Swarming and complex pattern formation in Paenibacillus vortex studied by imaging and tracking cells. BMC microbiology, 2008. 8: p. 36–36.

3. Kearns, D.B. and R. Losick, Swarming motility in undomesticated Bacillus subtilis. Mol Microbiol, 2003. 49(3): p. 581–90.

4. Kohler, T., L.K. Curty, F. Barja, C. van Delden, and J.C. Pechere, Swarming of Pseudomonas aeruginosa is dependent on cell-to-cell signaling and requires flagella and pili. J Bacteriol, 2000. 182(21): p. 5990–6.

5. Rauprich, O., M. Matsushita, C.J. Weijer, F. Siegert, S.E. Esipov, and J.A. Shapiro, Periodic phenomena in Proteus mirabilis swarm colony development. Journal of Bacteriology, 1996. 178(22): p. 6525.

6. Schuerle, S., A.P. Soleimany, T. Yeh, G. Anand, M. Häberli, H. Fleming, N. Mirkhani, F. Qiu, S. Hauert, and X. Wang, Synthetic and living micropropellers for convection-enhanced nanoparticle transport. Science advances, 2019. 5(4): p. eaav4803.

7. Li, S., R. Batra, D. Brown, H.-D. Chang, N. Ranganathan, C. Hoberman, D. Rus, and H. Lipson, Particle robotics based on statistical mechanics of loosely coupled components. Nature, 2019. 567(7748): p. 361–365.

8. Rubenstein, M., A. Cornejo, and R. Nagpal, Programmable self-assembly in a thousand-robot swarm. Science, 2014. 345(6198): p. 795–799.

9. Kearns, D.B., A field guide to bacterial swarming motility. Nature reviews. Microbiology, 2010. 8(9): p. 634–644.

10. Fujikawa, H. and M. Matsushita, Fractal Growth of Bacillus subtilis on Agar Plates. Journal of the Physical Society of Japan, 1989. 58(11): p. 3875–3878.

11. Prindle, A., P. Samayoa, I. Razinkov, T. Danino, L.S. Tsimring, and J. Hasty, A sensing array of radically coupled genetic ‘biopixels’. Nature, 2012. 481(7379): p. 39–44.

12. Payne, S., B. Li, Y. Cao, D. Schaeffer, M.D. Ryser, and L. You, Temporal control of self-organized pattern formation without morphogen gradients in bacteria. Molecular systems biology, 2013. 9(1): p. 697.

13. Santos-Moreno, J. and Y. Schaerli, Using Synthetic Biology to Engineer Spatial Patterns. Advanced Biosystems, 2019. 3(4): p. 1800280.

14. Basu, S., Y. Gerchman, C.H. Collins, F.H. Arnold, and R. Weiss, A synthetic multicellular system for programmed pattern formation. Nature, 2005. 434(7037): p. 1130–1134.

15. Liu, C., X. Fu, L. Liu, X. Ren, C.K.L. Chau, S. Li, L. Xiang, H. Zeng, G. Chen, L.-H. Tang, P. Lenz, X. Cui, W. Huang, T. Hwa, and J.-D. Huang, Sequential Establishment of Stripe Patterns in an Expanding Cell Population. Science, 2011. 334(6053): p. 238–241.

16. Curatolo, A., N. Zhou, Y. Zhao, C. Liu, A. Daerr, J. Tailleur, and J. Huang, Cooperative pattern formation in multi-component bacterial systems through reciprocal motility regulation. Nature Physics, 2020. 16(11): p. 1152–1157.

17. Sheth, R.U., S.S. Yim, F.L. Wu, and H.H. Wang, Multiplex recording of cellular events over time on CRISPR biological tape. Science, 2017. 358(6369): p. 1457–1461.

18. Perli, S.D., C.H. Cui, and T.K. Lu, Continuous genetic recording with self-targeting CRISPR-Cas in human cells. Science, 2016. 353(6304).

19. Frieda, K.L., J.M. Linton, S. Hormoz, J. Choi, K.-H.K. Chow, Z.S. Singer, M.W. Budde, M.B. Elowitz, and L. Cai, Synthetic recording and in situ readout of lineage information in single cells. Nature, 2017. 541(7635): p. 107–111.

20. Shipman, S.L., J. Nivala, J.D. Macklis, and G.M. Church, CRISPR–Cas encoding of a digital movie into the genomes of a population of living bacteria. Nature, 2017. 547(7663): p. 345–349.

21. Riglar, D.T., D.L. Richmond, L. Potvin-Trottier, A.A. Verdegaal, A.D. Naydich, S. Bakshi, E. Leoncini, L.G. Lyon, J. Paulsson, and P.A. Silver, Bacterial variability in the mammalian gut captured by a single-cell synthetic oscillator. Nature communications, 2019. 10(1): p. 1–12.

22. Schaffer, J.N. and M.M. Pearson, Proteus mirabilis and urinary tract infections, in Urinary Tract Infections: Molecular Pathogenesis and Clinical Management. 2017, ASM Press: Washington, DC. p. 383–433.

23. Cook, E.R. and N. Pederson, Uncertainty, emergence, and statistics in dendrochronology, in Dendroclimatology. 2011, Springer. p. 77–112.

24. Hauser, G., Uber Faulnisbakterien und deren Beziehung zur Septicamie. FGW Vogel, 1885.

25. Saak, C.C., K.A. Gibbs, and V.J. DiRita, The Self-Identity Protein IdsD Is Communicated between Cells in Swarming Proteus mirabilis Colonies. Journal of Bacteriology, 2016. 198(24): p. 3278–3286.

26. Fraser, G.M. and C. Hughes, Swarming motility. Curr Opin Microbiol, 1999. 2(6): p. 630–5.

27. Clemmer, K.M. and P.N. Rather, Regulation of flhDC expression in Proteus mirabilis. Research in Microbiology, 2007. 158(3): p. 295–302.

28. Howery, K.E., E. Simsek, M. Kim, and P.N. Rather, Positive autoregulation of the flhDC operon in Proteus mirabilis. Res Microbiol, 2018. 169(4-5): p. 199–204.

29. Pearson, M.M., D.A. Rasko, S.N. Smith, and H.L. Mobley, Transcriptome of swarming Proteus mirabilis. Infect Immun, 2010. 78(6): p. 2834–45.

30. Şimşek, E., E. Dawson, P.N. Rather, and M. Kim, Spatial regulation of cell motility and its fitness effect in a surface-attached bacterial community. The ISME journal, 2021: p. 1–8.

31. Armbruster, C.E., S.A. Hodges, and H.L. Mobley, Initiation of swarming motility by Proteus mirabilis occurs in response to specific cues present in urine and requires excess L-glutamine. Journal of bacteriology, 2013. 195(6): p. 1305–1319.

32. Little, K., J. Austerman, J. Zheng, and K.A. Gibbs, Cell shape and population migration are distinct steps of Proteus mirabilis swarming that are decoupled on high-percentage agar. Journal of bacteriology, 2019. 201(11): p. e00726–18.

33. Burall, L.S., J.M. Harro, X. Li, C.V. Lockatell, S.D. Himpsl, J.R. Hebel, D.E. Johnson, and H.L. Mobley, Proteus mirabilis genes that contribute to pathogenesis of urinary tract infection: identification of 25 signature-tagged mutants attenuated at least 100-fold. Infect Immun, 2004. 72(5): p. 2922–38.

34. Fraser, G.M., R.B. Furness, and C. Hughes, Swarming migration by Proteus and related bacteria. Prokaryotic Development, 1999: p. 379–401.

35. Huang, Z., X. Pan, N. Xu, and M. Guo, Bacterial chemotaxis coupling protein: Structure, function and diversity. Microbiological research, 2019. 219: p. 40–48.

36. Dufour, A., R.B. Furness, and C. Hughes, Novel genes that upregulate the Proteus mirabilis flhDC master operon controlling flagellar biogenesis and swarming. Molecular microbiology, 1998. 29(3): p. 741–751.

37. Hay, N.A., D.J. Tipper, D. Gygi, and C. Hughes, A nonswarming mutant of Proteus mirabilis lacks the Lrp global transcriptional regulator. Journal of Bacteriology, 1997. 179(15): p. 4741–4746.

38. Clemmer, K.M. and P.N. Rather, The Lon protease regulates swarming motility and virulence gene expression in Proteus mirabilis. Journal of Medical Microbiology, 2008. 57(8): p. 931–937.

39. Gygi, D., G. Fraser, A. Dufour, and C. Hughes, A motile but non-swarming mutant of Proteus mirabilis lacks FlgN, a facilitator of flagella filament assembly. Mol Microbiol, 1997. 25(3): p. 597–604.

40. Morgenstein, R.M. and P.N. Rather, Role of the Umo proteins and the Rcs phosphorelay in the swarming motility of the wild type and an O-antigen (waaL) mutant of Proteus mirabilis. Journal of bacteriology, 2012. 194(3): p. 669–676.

41. LeCun, Y., B. Boser, J.S. Denker, D. Henderson, R.E. Howard, W. Hubbard, and L.D. Jackel, Backpropagation Applied to Handwritten Zip Code Recognition. Neural Computation, 1989. 1(4): p. 541–551.

42. Russakovsky, O., J. Deng, H. Su, J. Krause, S. Satheesh, S. Ma, Z. Huang, A. Karpathy, A. Khosla, and M. Bernstein, Imagenet large scale visual recognition challenge. International journal of computer vision, 2015. 115(3): p. 211–252.

43. Jin, X. and I.H. Riedel-Kruse, Biofilm Lithography enables high-resolution cell patterning via optogenetic adhesin expression. Proceedings of the National Academy of Sciences, 2018. 115(14): p. 3698–3703.

44. Chen, A.Y., Z. Deng, A.N. Billings, U.O.S. Seker, Michelle Y. Lu, R.J. Citorik, B. Zakeri, and T.K. Lu, Synthesis and patterning of tunable multiscale materials with engineered cells. Nature Materials, 2014. 13: p. 515.

45. Huang, J., S. Liu, C. Zhang, X. Wang, J. Pu, F. Ba, S. Xue, H. Ye, T. Zhao, K. Li, Y. Wang, J. Zhang, L. Wang, C. Fan, T.K. Lu, and C. Zhong, Programmable and printable Bacillus subtilis biofilms as engineered living materials. Nature Chemical Biology, 2019. 15(1): p. 34–41.

46. Luo, N., S. Wang, and L. You, Synthetic pattern formation. Biochemistry, 2019. 58(11): p. 1478–1483.

47. Nasip, Ö.F. and K. Zengin. Deep Learning Based Bacteria Classification. in 2018 2nd International Symposium on Multidisciplinary Studies and Innovative Technologies (ISMSIT). 2018. IEEE.

48. Casado-García, Á., G. Chichón, C. Domínguez, M. García-Domínguez, J. Heras, A. Inés, M. López, E. Mata, V. Pascual, and Y. Sáenz, MotilityJ: An open-source tool for the classification and segmentation of bacteria on motility images. Comput Biol Med, 2021. 136: p. 104673.

49. Wang, H., H.C. Koydemir, Y. Qiu, B. Bai, Y. Zhang, Y. Jin, S. Tok, E.C. Yilmaz, E. Gumustekin, and Y. Rivenson, Early-detection and classification of live bacteria using time-lapse coherent imaging and deep learning. arXiv preprint arXiv:2001.10695, 2020.

50. Jeckel, H., E. Jelli, R. Hartmann, P.K. Singh, R. Mok, J.F. Totz, L. Vidakovic, B. Eckhardt, J. Dunkel, and K. Drescher, Learning the space-time phase diagram of bacterial swarm expansion. Proceedings of the National Academy of Sciences, 2019. 116(5): p. 1489–1494.

51. Lugagne, J.-B., H. Lin, and M.J. Dunlop, DeLTA: Automated cell segmentation, tracking, and lineage reconstruction using deep learning. PLoS computational biology, 2020. 16(4): p. e1007673.

52. Doshi, A., M. Shaw, R. Tonea, R. Minyety, S. Moon, A. Laine, J. Guo, and T. Danino, A deep learning pipeline for segmentation of *Proteus mirabilis* colony patterns. bioRxiv, 2022: p. 2022.01.17.475672.

53. Dietrich, L.E., T.K. Teal, A. Price-Whelan, and D.K. Newman, Redox-active antibiotics control gene expression and community behavior in divergent bacteria. Science, 2008. 321(5893): p. 1203–1206.

54. Minogue, T., H. Daligault, K. Davenport, K. Bishop-Lilly, D. Bruce, P. Chain, S. Coyne, O. Chertkov, T. Freitas, and K. Frey, Draft genome assemblies of Proteus mirabilis ATCC 7002 and Proteus vulgaris ATCC 49132. Genome announcements, 2014. 2(5).

55. Pearson, M.M., Methods for Studying Swarming and Swimming Motility, in Proteus mirabilis: Methods and Protocols, M.M. Pearson, Editor. 2019, Springer New York: New York, NY. p. 15–25.

56. Tol, P., Colour Schemes. 2021, SRON: SRON/EPS/TN/09-002.

57. Kutschera, A. lightM. Available from: https://github.com/vektorious/lightM.

58. Levin-Reisman, I., O. Gefen, O. Fridman, I. Ronin, D. Shwa, H. Sheftel, and N.Q. Balaban, Automated imaging with ScanLag reveals previously undetectable bacterial growth phenotypes. Nature Methods, 2010. 7(9): p. 737–739.

59. Bär, J., M. Boumasmoud, R.D. Kouyos, A.S. Zinkernagel, and C. Vulin, Efficient microbial colony growth dynamics quantification with ColTapp, an automated image analysis application. Scientific reports, 2020. 10(1): p. 1–15.

60. Hartmann, R., H. Jeckel, E. Jelli, P.K. Singh, S. Vaidya, M. Bayer, D.K. Rode, L. Vidakovic, F. Díaz-Pascual, and J.C. Fong, Quantitative image analysis of microbial communities with BiofilmQ. Nature microbiology, 2021. 6(2): p. 151–156.

61. Pavlov, L. moving_polyfit. 2021 August 30, 2021]; Available from: https://www.mathworks.com/matlabcentral/fileexchange/86503-moving_polyfit.

62. Hand, D.J. and R.J. Till, A Simple Generalisation of the Area Under the ROC Curve for Multiple Class Classification Problems. Machine Learning, 2001. 45(2): p. 171–186.

63. Russell, B.C., A. Torralba, K.P. Murphy, and W.T. Freeman, LabelMe: a database and web-based tool for image annotation. International journal of computer vision, 2008. 77(1-3): p. 157–173.

64. Sundararajan, M., A. Taly, and Q. Yan. Axiomatic attribution for deep networks. in International Conference on Machine Learning. 2017. PMLR.

65. He, K., X. Zhang, S. Ren, and J. Sun, Deep Residual Learning for Image Recognition. 2016 IEEE Conference on Computer Vision and Pattern Recognition (CVPR), 2016: p. 770–778.

66. Szegedy, C., W. Liu, Y. Jia, P. Sermanet, S.E. Reed, D. Anguelov, D. Erhan, V. Vanhoucke, and A. Rabinovich, Going deeper with convolutions. 2015 IEEE Conference on Computer Vision and Pattern Recognition (CVPR), 2015: p. 1–9.

67. Szegedy, C., V. Vanhoucke, S. Ioffe, J. Shlens, and Z. Wojna, Rethinking the Inception Architecture for Computer Vision. 2016 IEEE Conference on Computer Vision and Pattern Recognition (CVPR), 2016: p. 2818–2826.

68. Ronneberger, O., P. Fischer, and T. Brox. U-Net: Convolutional Networks for Biomedical Image Segmentation. 2015. Cham: Springer International Publishing.

69. Iglovikov, V. and A. Shvets, Ternausnet: U-net with vgg11 encoder pre-trained on imagenet for image segmentation. arXiv preprint arXiv:1801.05746, 2018.

70. Simonyan, K. and A. Zisserman, Very deep convolutional networks for large-scale image recognition. arXiv preprint arXiv:1409.1556, 2014.

