## Supplemental Figures and Tables for "Engineered bacterial swarm patterns as spatial records of environmental inputs"

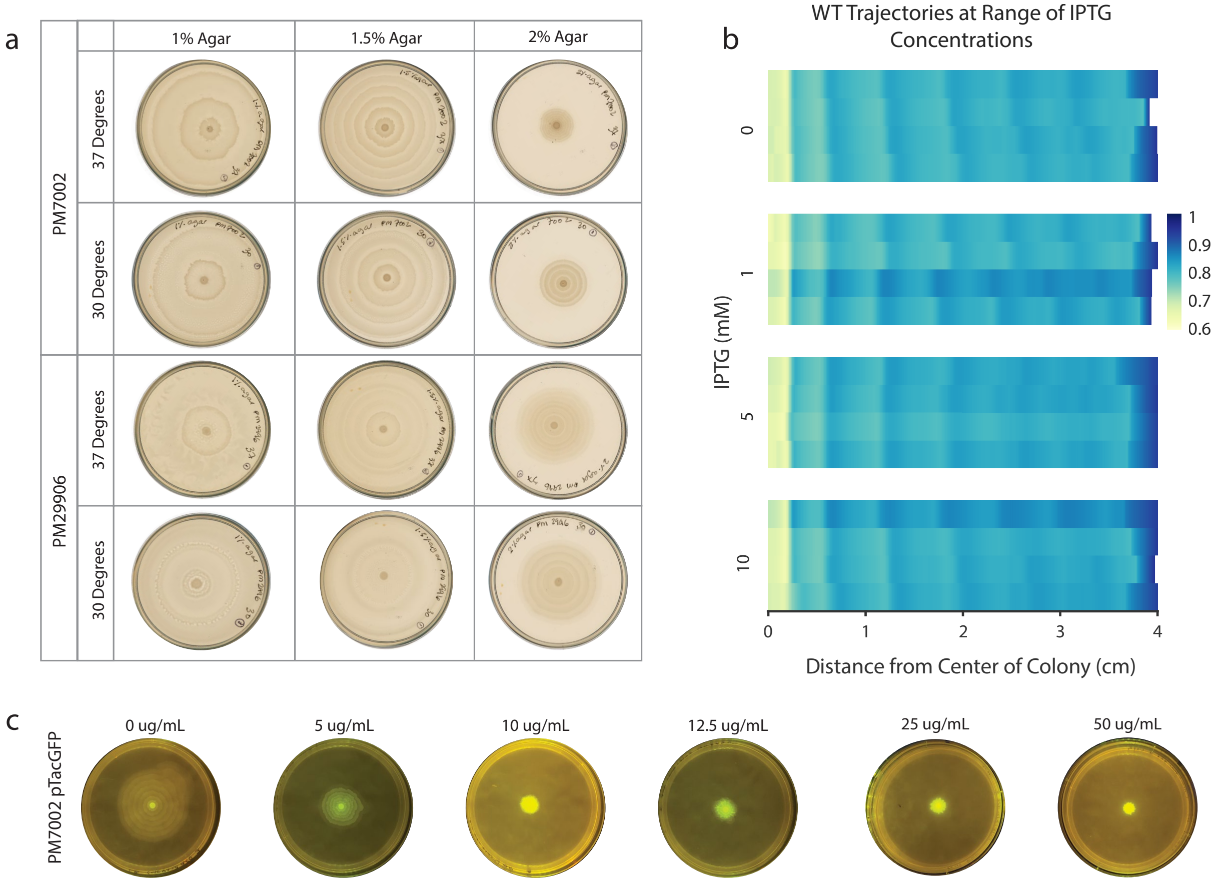


**Figure S1: Selection of Conditions. a.** Representative colony images for different *P. mirabilis* strains, temperatures, and agar hardness after 24 hours of growth. **b.** Heatmaps of radially averaged profiles of PM7002 colonies at a range of IPTG concentrations (each profile represents an individual colony). Light green represents lowest pixel intensity (highest colony density), a.u. **c.** PM7002 with pTacGFP plasmid was grown at a range of kanamycin concentrations for 24 hours. Representative images of fluorescence taken under blue light transilluminator are shown.

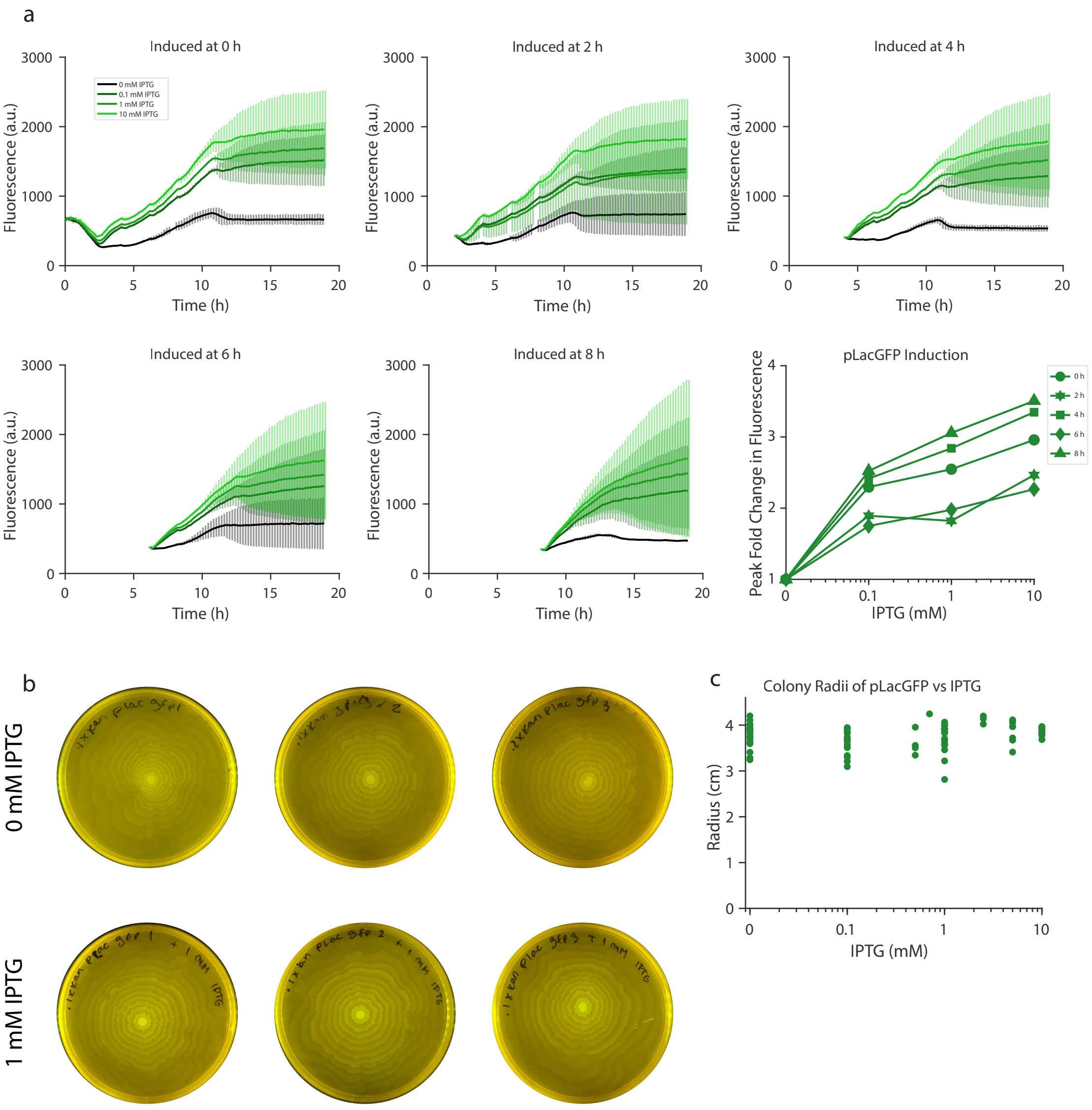


**Figure S2: Induction of pLac promoter in *P. mirabilis*. a.** Fresh overnight cultures of PM7002 with pLacGFP plasmid were subcultured with 50 ug/mL kanamycin, grown for 0-8 hours, and induced with a range of IPTG concentrations. The mean GFP intensity (arbitrary units) after subtracting background fluorescence and normalizing to OD is plotted for each group; error bars indicate standard deviation (n = 3 for each). Bottom right panel: Peak fold change in fluorescence from uninduced (0 mM IPTG) groups is plotted for each group induced at a different time after subculturing (mean of three replicates, individual data shown in **a**). Maximal expression of around 3-fold was consistent with literature[1]. **b.** Images of PM7002 pLacGFP strain plates grown with either 0 or 1 mM IPTG, imaged after 24 hours under blue-light transilluminator. **c.** 24-hour colony radii of pLacGFP strain at various concentrations of IPTG; each dot represents a plate. Plate images are from different days.

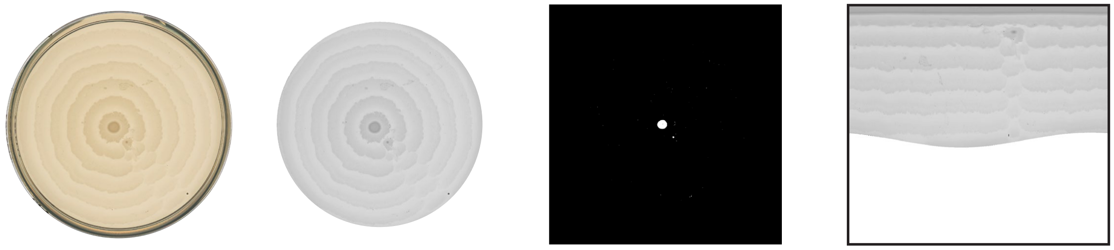

**Figure S3: Image processing.** Petri dishes were scanned at high resolution. The Petri rim was identified and cropped out using MATLAB functions. Images were thresholded to show only the colony inoculum, and the center point was identified using MATLAB functions. Images were converted from Cartesian to polar coordinates with interpolation, and the flattened images were used for subsequent analysis. See Methods for details.

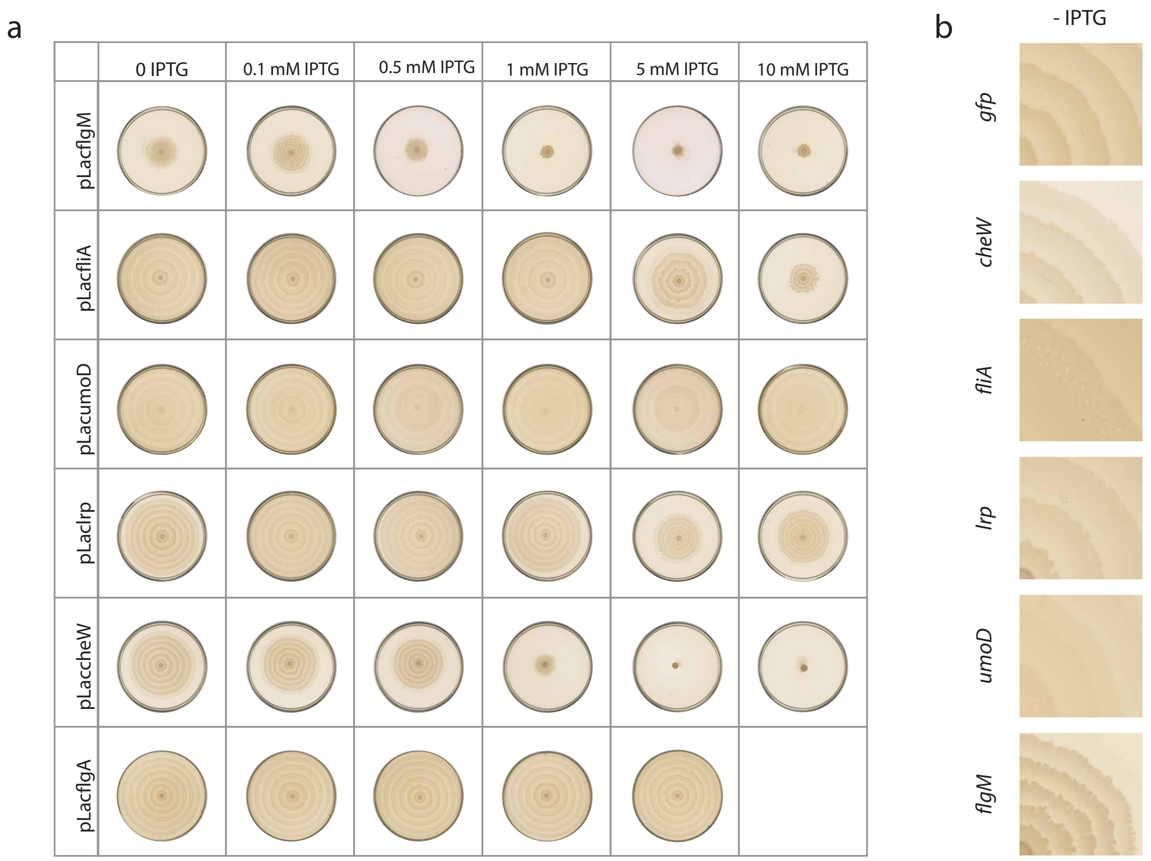


**Figure S4: Range of patterns formed. a.** PM7002 with indicated inducible swarm plasmids was grown for 24 hours at a range of IPTG concentration. Representative images of three replicates are shown. **b.** Closeups of patterns formed at 0 mM IPTG.

a

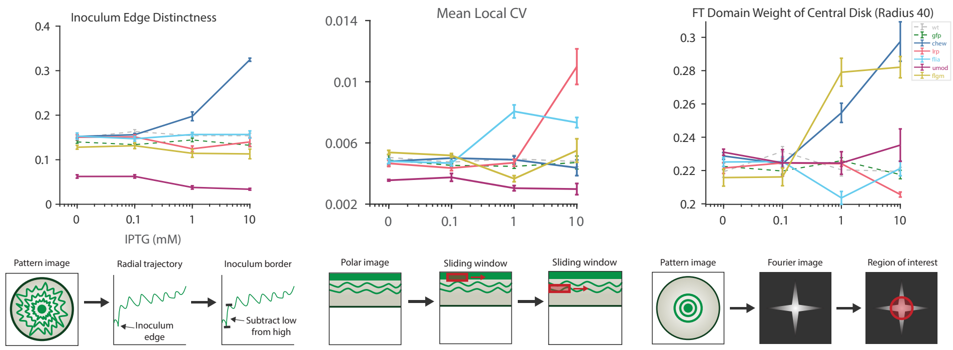


b

| Strain | Colony Sector Mean Intensity | p | Colony Sector Across-Ring CV | p | Colony Sector Local CV | p | Colony Sector Min Intensity | p | Colony Sector Area | p | AUC |
|---|---|---|---|---|---|---|---|---|---|---|---|
| WT | -91.9 | 1.5e-7 | 101.6 | 0.11 | 94.4 | 0.72 | | | | | 0.667 |
| gfp | -17.9 | 0.025 | 24.0 | 0.026 | -314.57 | 1.9e-5 | -2.0 | 0.32 | -5.3 | 0.001 | 0.60 |
| lrp | 30.3 | 0.26 | 176.3 | 6.7e-2 | -2919 | 8.5e-6 | -55.7 | 5.3e-6 | -26.0 | 6.1e-2 | 0.965 |
| cheW | -9.3 | 0.026 | -27.5 | 8.5e-2 | | | -4.6 | 0.13 | 13.6 | 6.1e-2 | 0.90 |
| flgM | | | -71.6 | 1.8e-5 | 193.9 | 6.6e-4 | 6.8 | 6.8e-3 | 10.5 | 2.8e-8 | 0.811 |
| fliA | | | 443.7 | 2.4e-3 | -2324 | 1.3e-7 | 62.2 | 7.9e-3 | 77.6 | 0.51 | 0.959 |

| Gene | Inoculum Max Intensity | p | Inoculum Mean Intensity | p | Inoculum Variance | p | AUC |
|---|---|---|---|---|---|---|---|
| umoD | -36.6 | 0.721 | -205.0 | 0.046 | 31131 | 0.065 | 0.921 |

c

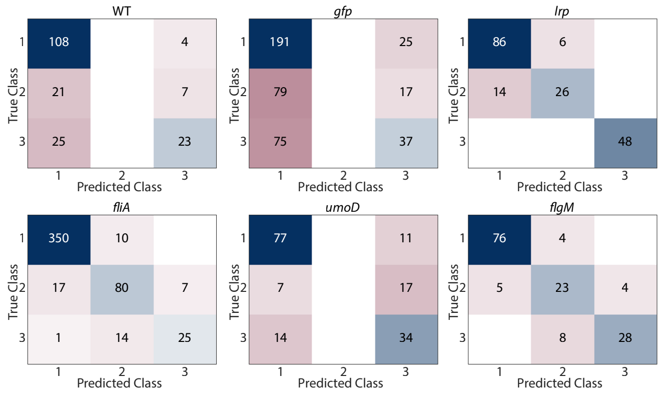


**Figure S5: Quantification of engineered colony patterns. a.** Quantification of aspects of colony patterns of engineered strains at increasing IPTG concentrations, measured as in Figure 2e-g and as shown in the schematics below each graph. All strains had at least n = 3 plates measured at each IPTG concentration. Error bars represent standard error of the mean. **b.** Multinomial regression model performance on engineered strains. Each flattened image was divided into four sectors (each 250 columns wide) to augment number of measurements available and allow model to converge. Measurements listed in **b** were obtained on each sector and mnrfit function was used to obtain a best fit model for classifying sets of measurements into bins 1: 0-0.9 mM IPTG, 2: 1-5 mM IPTG, 3: 5-10 mM IPTG. The table shows each variable's coefficient and p-value of the coefficient, both returned by mnrfit for the best fit model of each strain, along with the AUC of the model. **c.** Confusion matrices of the fitted models' performance on each dataset; numbers on the matrices represent numbers of plates.

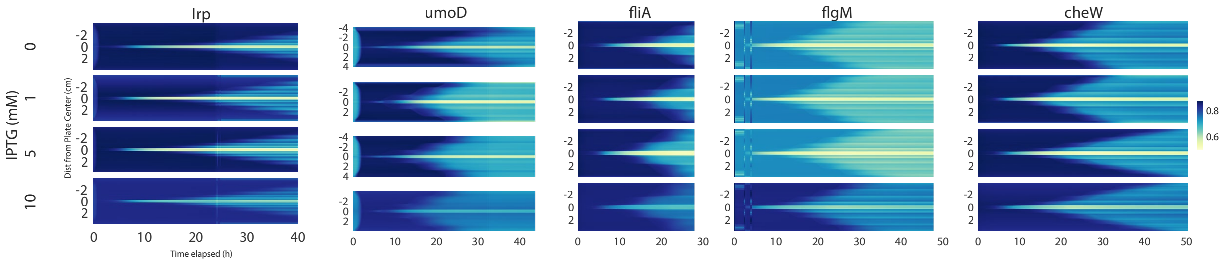


**Figure S6: Tuning dynamics of *P. mirabilis* pattern formation by strain.** Time-lapse growth of various engineered *P. mirabilis* strains at varying IPTG concentrations are displayed as heatmaps. Each heatmap represents one colony, with blue representing highest pixel intensity (lightest/less dense areas of colony) and yellow representing densest colony portions. The colony front can be identified as a faint light-colored line emanating outwards from the initial center inoculum (brightest horizontal bar across each heatmap).

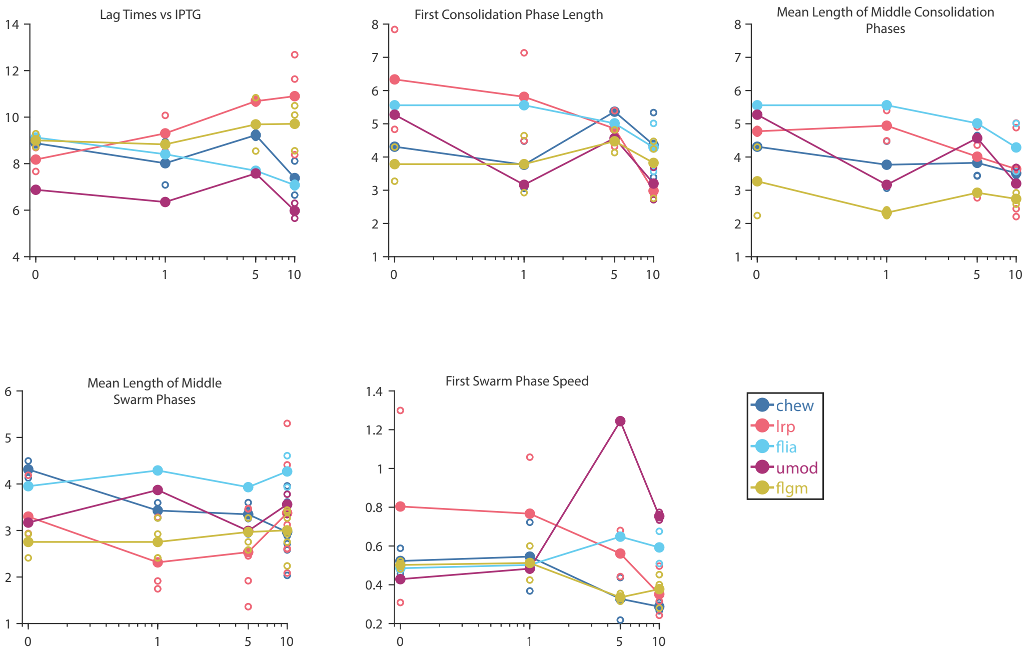


**Figure S7: Dynamics of pattern formation.** Plots of dynamic characteristics of engineered strains vs IPTG concentration, measured from one to two time-lapses per condition per strain. Individual dots represent individual time-lapses (lag times) or individual phases in all time-lapses (consolidation and swarm phase lengths, swarm speeds). Lines represent the mean of these individual measurements. Middle swarm or consolidation phase lengths were determined by discarding the measurements of the first and last of these phases for each time-lapse, or discarding the last phase if the time-lapse had only two of the given phase.

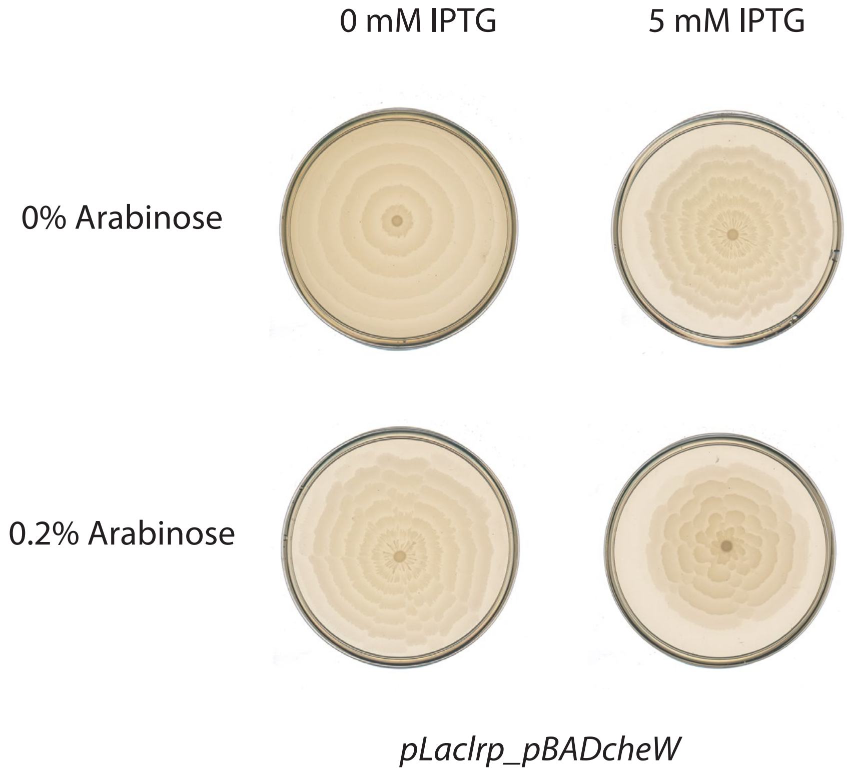


**Figure S8: *lrp* and *cheW* combination strain.** Representative images of the *pLaclrp_pBadcheW* strain at given concentrations of arabinose and IPTG.

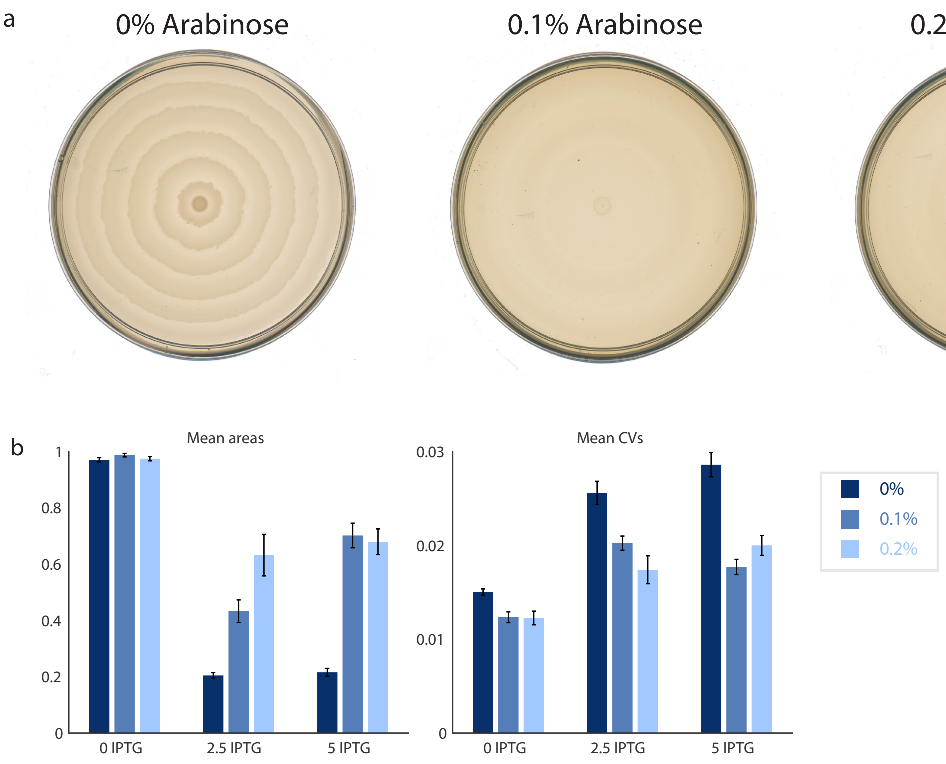


**Figure S9: Building and characterizing *cheWumoD* combination strain. a.** *pBADumoD* control. Representative images of colonies after 24 hours of growth on 1.5% agar with indicated arabinose concentrations are shown. **b.** Plots of the mean area (percent of plate) and average radial coefficient of variation of *cheWumoD* patterns shown in **Fig. 4d-e**; error bars represent standard error of the mean. At 0 mM IPTG, number of plates = 41, 35, 31 for 0, 0.1, 0.2% arabinose respectively; at 2.5 mM IPTG, number of plates = 23, 22, 23 for 0, 0.1, 0.2% arabinose respectively; at 5 mM IPTG, number of plates = 27, 14, 26 for 0, 0.1, 0.2% arabinose respectively. For colony area, comparing 0.1% to 0.2% arabinose, $p = 0.0145$ for two-sample t-test at 2.5 mM IPTG, but at 0 IPTG $p = 0.2235$ and at 5 IPTG $p = 0.7$.

a

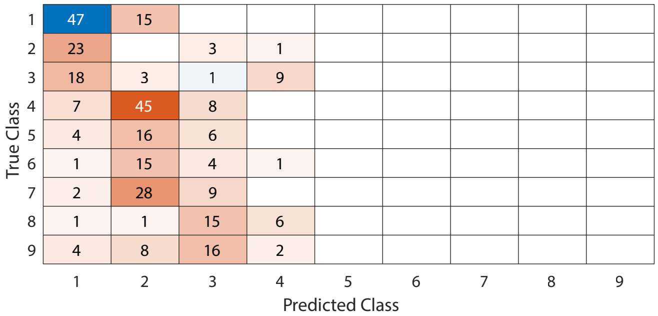


b

| | Weights |
|---|---|
| Col. Radius (Pixels) | -0.0223 |
| Col. Radius (cm) | 3.7914 |
| Reached Edge of Plate | 2.5727 |
| Mean Colony Intensity | -11.2744 |
| Mean Sliding CV | -48.8359 |
| Mean Inoculum Intensity | -18.9936 |
| Max Inoculum Intensity | 18.8092 |

**Figure S10: Multinomial regression for dual input strain. a.** Multinomial regression model performance on dual input strain measurements. The mnrfit function was used to obtain a best fit model for classifying sets of measurements into nine bins: 1: 0-1 mM IPTG, 0-0.1% arabinose; 2: 1-5 mM IPTG, 0-0.1% arabinose; 3: 5-10 mM IPTG, 0-0.1% arabinose; 4: 0-1 mM IPTG, 0.1-0.19% arabinose; 5: 1-5 mM IPTG, 0.1-0.19% arabinose; 6: 5-10 mM IPTG, 0.1-0.19% arabinose; 7: 0-1 mM IPTG, 0.2% arabinose; 8: 1-5 mM IPTG, 0.2% arabinose; 9: 5-10 mM IPTG, 0.2% arabinose. The confusion matrix was then plotted; numbers on the matrix represent numbers of plates. **b.** Weights on each variable used for the best fit model.

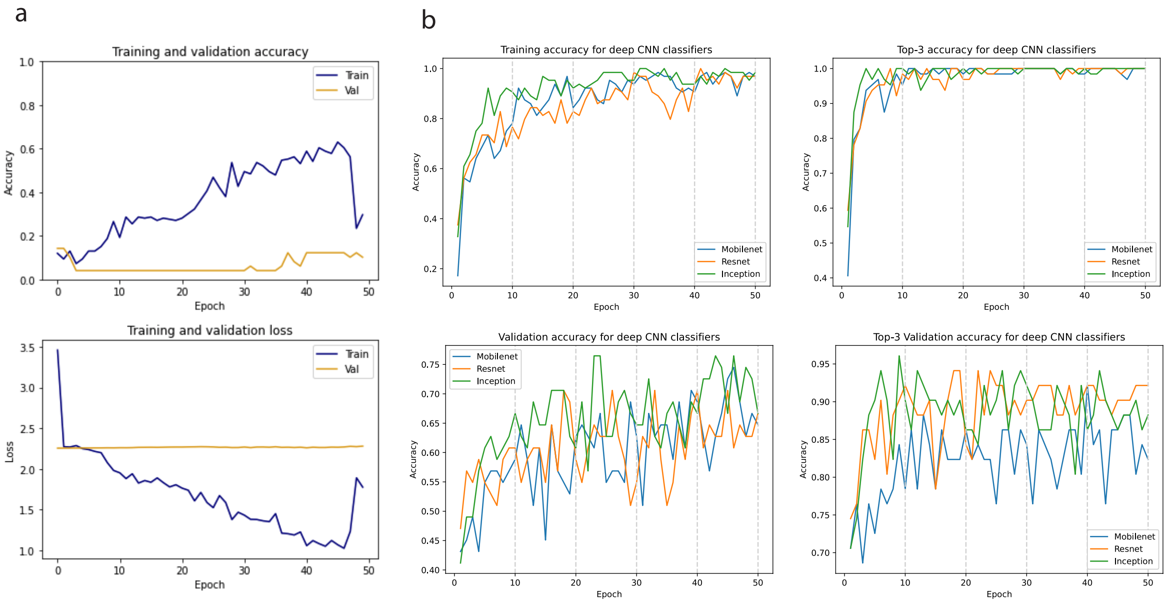


c

| Model | Mean Accuracy |
|---|---|
| kNN classifier | 0.197674 |
| SVC with linear kernel | 0.312302 |
| SVC with RBF kernel | 0.314250 |
| VGG19 | 0.726149 |
| ResNet18V2 | 0.767558 |
| **MobileNetV2** | **0.824543** |
| **ResNet50V2** | **0.833333** |
| **InceptionV3** | **0.841954** |

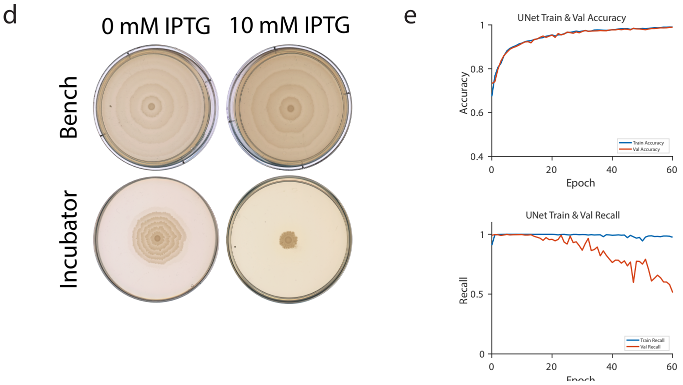


**Figure S11: Performance of CNNs on classification and segmentation tasks. a.** Training and validation accuracy and loss for a CNN model which had three convolutional/max pooling blocks, trained on the dataset of images of the combination strain at various IPTG and arabinose conditions (same dataset as in **Fig. 4f**). **b.** Fine tuning of three architectures pre-trained on ImageNet weights; right hand panels represent models' ability to identify the correct image class within its top three predicted classes. **c.** Mean accuracy on a separated test dataset of combination strain images for each of the fine-tuned pre-trained architectures (80/20 split of dataset shown in **Fig. 4f**). **d.** Representative images of *pLacflgM* strain pattern at the given conditions. The difference between incubator and benchtop patterns is most extreme at 10 mM IPTG. **e.** Training and validation accuracy and recall of the U-Net model with VGG-11 encoder, trained and validated on 17 (13 training, 4 validation) images and evaluated on 4 test images.

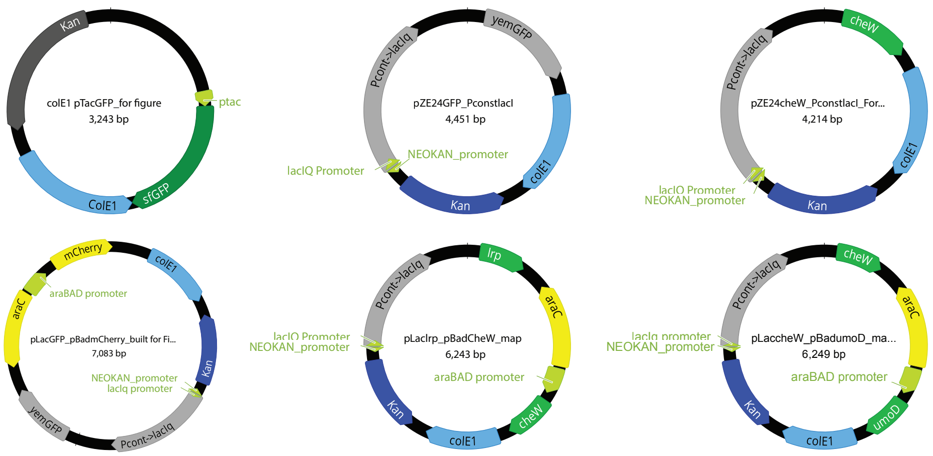


**Figure S12: Plasmids used.**

**Table S1 Bacterial Strains and Plasmids**

| **Label** | **Host Strain** | **Plasmid** |
|---|---|---|
| AD275 | *P. mirabilis* (PM7002) | colE1pTacGFP |
| AD283 | *P. mirabilis* (PM7002) | pze24pConstlacIQ |
| AD300 | *P. mirabilis* (PM7002) | pLacflgM |
| AD308 | *P. mirabilis* (PM7002) | pLaclrp |
| AD350 | *P. mirabilis* (PM7002) | pLacflgA |
| AD375 | *P. mirabilis* (PM7002) | pLacumoD |
| AD379 | *P. mirabilis* (PM7002) | pLacfliA |
| AD391 | *P. mirabilis* (PM7002) | pLaccheW |
| AD356 | *P. mirabilis* (PM7002) | pBadmCherry |
| AD432 | *P. mirabilis* (PM7002) | placcheW-pBadumoD |
| AD434 | *P. mirabilis* (PM7002) | placlrp-BadcheW |

**Table S2 Swarm Gene Primers for Gibson Assembly**

| Primer Name | Sequence |
|---|---|
| fliA_r | CGTTTTATTTGATGCCTCTAGCACGCGTCTATTTCTCTGGGTTTAAGCG |
| fliA_f | CACACAGAATTCATTAAAGAGGAGAAAGGTACCGTG AGT GAT TTG TAT ACC GCC |
| lrp_f | CACACAGAATTCATTAAAGAGGAGAAAGGTACCATGATTGATAATAAAAAACGTCCGGG |
| lrp_r | CGTTTTATTTGATGCCTCTAGCACGCGTTTAGCGTGTTTTAATCACTAGGC |
| umoD_f | CACACAGAATTCATTAAAGAGGAGAAAGGTACCGTGAGTGTTGATAGCAAAAAGTTC |
| umoD_r | CGTTTTATTTGATGCCTCTAGCACGCGTTTACTCTTTACGGCAAGGAATATTTTC |
| flgM_f | CACACAGAATTCATTAAAGAGGAGAAAGGTACCATGAGTATTGAACGCACAAATCC |
| flgM_r | CGTTTTATTTGATGCCTCTAGCACGCGTTTATTTAGTTATGCTTTCAGCAGC |
| flgA_f | CACACAGAATTCATTAAAGAGGAGAAAGGTACCATGTCTGTTAGAACGTTTATCGG |
| flgA_r | CGTTTTATTTGATGCCTCTAGCACGCGTTTACAAGGGGATACGTACAGAG |
| cheW_r | CGTTTTATTTGATGCCTCTAGCACGCGTTTATTTAGCCATAATTGTTGCG |
| cheW_f | CACACAGAATTCATTAAAGAGGAGAAAGGTACCATGGCTGCGGAACATTTC |

**Movie S1**

Time-lapse movie of engineered strains and *gfp* control, with images captured every 10 minutes as described in Methods. All plates contained 10 mM IPTG. Images were downsampled 0.25x in movie to reduce file size but used at full size during analysis.

References

1. Baumberg, S. and M. Roberts, *Anomalous expression of the E. coli lac operon in Proteus mirabilis.* Molecular and General Genetics MGG, 1984. **198**(1): p. 166-171.
